# Organotchi: an open-source tool for the automated segmentation, morphological characterization, and longitudinal tracking of organoids

**DOI:** 10.64898/2026.11.04.764279

**Authors:** Lízia Branco, Thomas Lautenschlager, Daria Andreeva, Zeynep Yentür, Francesco Castagnetti, Theresa Kagermeier, Isabel Potthof, Pierre Collignon, Kensington Potter Stephens, Julia Matilainen, Ralf Mikut, Simone Mayer

## Abstract

Neural organoids are three-dimensional stem cell-derived models widely used to study brain development and model disease. Brightfield imaging offers a non-invasive readout of their growth and morphology, and recent advances have made automated organoid segmentation from such images increasingly attainable. Yet existing tools largely leave morphological characterization, quality assessment, and comparisons between conditions to manual or *ad hoc* analysis. Here we present Organotchi, an open-source tool that integrates automated segmentation with longitudinal morphological analysis of neural organoids. Organotchi quantifies ten morphological features, including area, perimeter, and eccentricity, and tracks cystic structures over time, reports variability within and between batches, and flags outlier organoids and segmentation failures, supporting objective organoid characterization and enhancing downstream analysis. As a proof of concept, we apply Organotchi to disease modelling, pharmacological perturbation, and protocol optimization in neural organoids. One use case assesses drug response in breast cancer spheroids, demonstrating its use beyond its development domain.

## Introduction

Neural organoids are three-dimensional (3D) culture systems generated from pluripotent stem cells (PSCs) or adult stem cells (Kim *et al*, 2020). Because they reproduce defined aspects of the cytoarchitecture, cellular composition, and developmental trajectory of the human brain (Clevers, 2016; Lancaster et al, 2013), they are widely used to investigate the effects of genetic or environmental perturbations on neurodevelopment (Pașca, 2018; Eichmüller & Knoblich, 2022; Kelley & Pașca, 2022). More broadly, organoid models are increasingly expected to contribute to preclinical and regulatory decisions (European Commission, 2026; U.S. Food and Drug Administration (FDA), 2025), which is driving the field towards greater standardization – for example through reporting frameworks and morphology-based quality control (Pașca *et al*, 2022; Castiglione *et al*, 2025; Kegeles *et al*, 2020; Marsee *et al*, 2021) – and higher throughput – through the automation of organoid generation and experimental readouts (Renner *et al*, 2020; Czerniecki *et al*, 2018; Brandenberg *et al*, 2020; Xu *et al*, 2025).

Importantly, the same properties that give these models their physiological relevance, namely their capacity for self-organization and the flexibility of the starting cell material, are also intrinsic sources of variability. Differences between human PSC lines and between clones of the same line consistently give rise to some level of heterogeneity, as do differences between batches of the same protocol (Boerstler *et al*, 2025; Strano *et al*, 2020; Øhlenschlæger *et al*, 2026; Branco *et al*, 2026). Additionally, stochastic events during neural induction can produce organoids enriched in off-target, non-neural tissue (Jensen & Little, 2023; Bhaduri *et al*, 2020). Protocol development and refinement is therefore in large part an exercise in identifying, quantifying, and constraining these sources of variability across organoid differentiation and maturation. At the same time, how confidently a biological effect can be distinguished from background heterogeneity depends on how well that heterogeneity has been measured and tracked over time, whether it manifests in overall size and shape, or in cellular composition and cytoarchitecture, or at the molecular and functional level.

Longitudinal analysis of organoid shape, here referred to as morphology, emerges as an informative way to measure and track this heterogeneity, usually by brightfield or phase-contrast microscopy – a label-free, inexpensive modality compatible with routine culture handling. Other non-invasive imaging modalities, such as high-field magnetic resonance imaging, are also possible (Deininger *et al*, 2023), but at substantially lower spatial resolution and considerably greater experimental effort. As a quality readout, external morphology can be an indicator of the ongoing differentiation (Chiaradia et al, 2023) and reveal the presence of off-target lineages (Castiglione *et al*, 2025). Several groups have therefore implemented morphology-based, non-destructive selection and scoring schemes as a quality-control step for cerebral organoid cultures (Castiglione et al, 2025; Boerstler et al, 2025; Chiaradia et al, 2026). Additionally, longitudinal analysis of organoid external morphology can also serve as a phenotypic readout. For example, organoid models of Zika virus infection (Dang et al, 2016; Garcez et al, 2016; Watanabe et al, 2017) and prenatal valproic acid exposure (Yentür et al, 2026; Cui et al, 2020) have reported reduced organoid size, while PTEN deletion resulted in substantially larger and folded cerebral organoids (Li *et al*, 2017).

Analysis of organoid external morphology can be accomplished either manually, against expert-held qualitative criteria, or through automated alternatives. Manual assessment may suffice for small experiments but it is slow, operator-dependent, difficult to audit, and incompatible with multi-batch longitudinal designs. Automated tools fall into several broad categories. Machine learning-free pipelines identify organoids using intensity thresholds and simple shape filters, are transparent, and require no training data (Borten et al, 2018; Wells et al, 2026; Stirling et al, 2021). Deep-learning tools trained specifically on organoid images achieve high segmentation accuracy (Powell *et al*, 2022; Qin *et al*, 2025; Kassis *et al*, 2019; Matthews *et al*, 2022; Park *et al*, 2023; Lefferts *et al*, 2024; Schröter *et al*, 2024), but their performance often drops when applied outside of their development domain. More recently, generalist segmentation models, trained on large and diverse image collections, have reduced this limitation. These include models developed for cellular segmentation (Stringer *et al*, 2021), promptable foundation models (Kirillov *et al*, 2023; Archit *et al*, 2025; Pachitariu *et al*, 2025), and pipelines that pair a detector with prompt-guided segmentation to handle overlapping instances (Deng et al, 2026). As reliable segmentation across organoid types becomes increasingly attainable, the main bottleneck has shifted from identifying and segmenting organoids to determining which morphological features to measure and how these measurements can inform standardization, quality control, and phenotypic characterization. To our knowledge, no existing tool yet addresses this by integrating experiment-level quality control throughout the culture period with a comprehensive morphological analysis pipeline, so that intra- and inter-batch variability and differences between experimental conditions can be assessed rapidly and with minimal user input.

To address this gap, we developed Organotchi, an open-source tool for the automated segmentation, morphological characterization, and longitudinal tracking of organoids from brightfield images. Although Organotchi was designed primarily for brain organoid research, we show that it also generalizes to other 3D culture models. Organotchi integrates two connected but independent frameworks (Figure 1): one converting raw acquisitions into per-organoid masks, and one extracting external morphological features, such as size and shape, from those masks and comparing them within and between batches and/or experimental conditions. Either framework can be used without the other, so that masks produced manually or by an external tool can be passed directly to the analysis framework, and the segmentation output of Organotchi can be exported into the characterization pipeline of the user. To make longitudinal morphological analysis robust and informative, as well as accessible and adaptable across the full organoid culture period, each step of the workflow allows an explicit user choice. Segmentation, for example, may proceed through a classical or a deep-learning-based (Deng *et al*, 2026; Xie *et al*, 2021) route, the latter offering two models built on different architectures. These routes were selected upon the benchmarking of eight previously reported segmentation approaches. Beyond size and shape, Organotchi can optionally partition each organoid into lighter and darker regions, providing a proxy for cystic area that can be quantified and tracked over time, whether cysts are an indicator of off-target differentiation (Ikeda *et al*, 2024), or an expected feature, such as in choroid plexus organoids (Pellegrini *et al*, 2020). The pipeline also reports on its own performance, including undetected organoids and observations flagged as outliers, which can optionally be excluded from subsequent analysis. This supports objective organoid characterization and more reliable downstream analysis.

**Figure 1.**
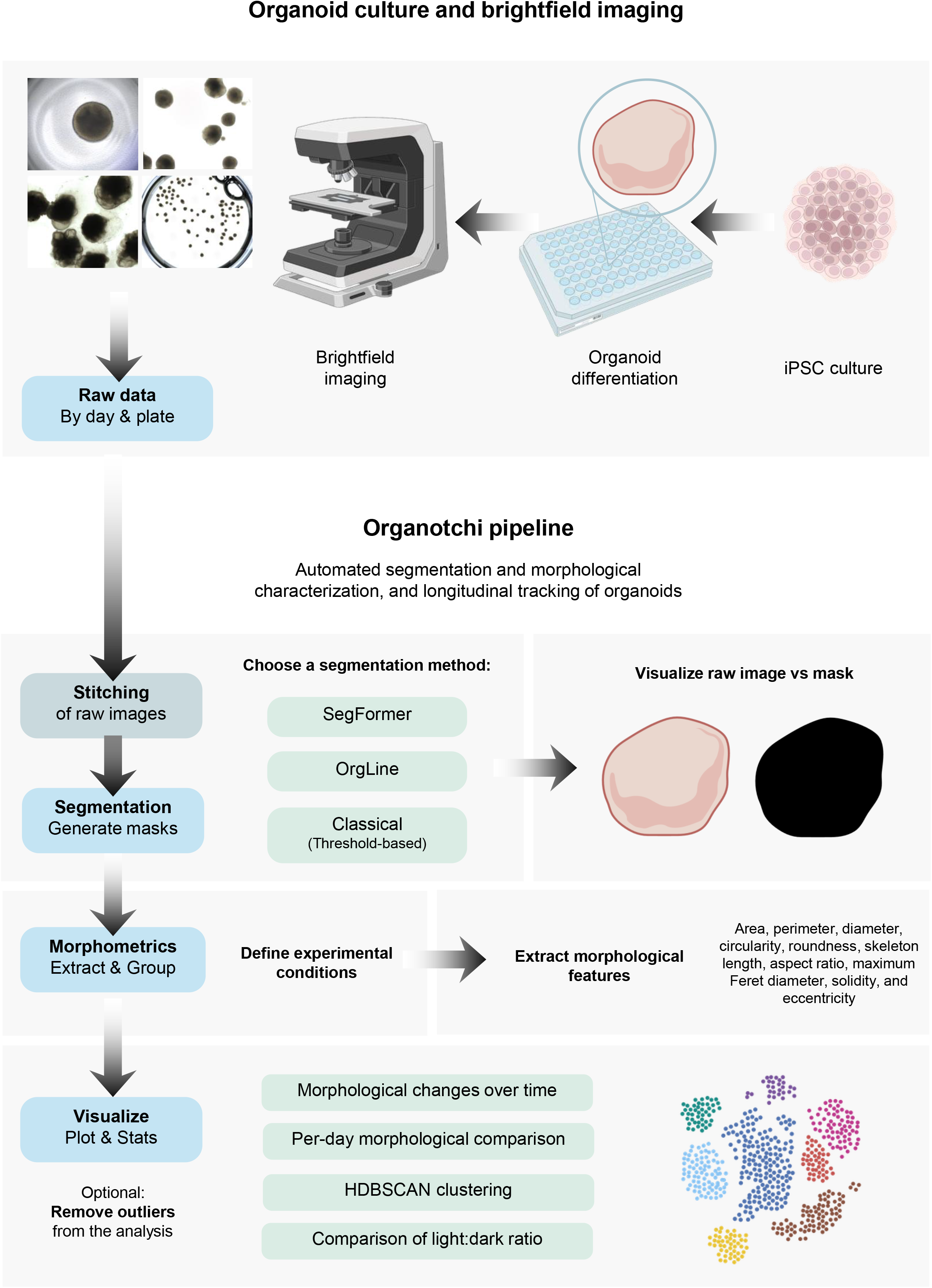
Organotchi pipeline workflow. *Segmentation framework:* Following image acquisition, the user provides the path to the folder containing the raw data for a given project, organized by time point (day) and imaged plate. Image processing parameters are then specified (for example, whether multiple tiles per well require stitching, and the pixel-to-micron calibration), and a segmentation method is chosen, which may differ between time windows and plates. The user can additionally choose whether light and dark sub-regions within the organoids are to be detected. Once segmentation has run, the resulting masks can be inspected side by side with the raw (stitched) images within the tool. *Morphometrics framework:* The user selects the morphological features to be extracted from the masks and defines the experimental groups into which the imaged plates fall. Results can likewise be visualized within the tool, through longitudinal plots, per-day comparisons of morphological features and their rates of change across groups (with pairwise testing), dimensionality reduction on non-redundant features using UMAP, and HDBSCAN clustering, with analysis of cluster composition by condition and of the morphological features characterizing each cluster.

As a proof of concept, we apply Organotchi to four independent experiments. Case studies 1, 2 and 4 involved neural organoids and covered different imaging settings, plating strategies, and organoid regional identities and morphologies, addressing disease modelling, drug perturbation, and protocol optimization. Case study 3 consisted on a drug treatment of breast cancer spheroids and tested Organotchi outside its development domain. These experiments draw on both published and previously unreported imaging datasets, the latter comprising more than 3,500 image-mask pairs, which we release alongside Organotchi.

## Results

### Selection of segmentation approaches to include within Organotchi

Organotchi was designed to support objective organoid characterization by integrating, within a single tool, the segmentation of brightfield organoid images, the morphological analysis of the resulting masks, and the extraction of information relevant to both quality control (including within-group variability and outlier detection) and phenotypic characterization across experimental conditions. To this end, Organotchi comprises two independent but connected frameworks, one dedicated to image segmentation and the other to morphological analysis (Figure 1).

As all downstream measurements rely on mask quality, we first benchmarked previously reported segmentation approaches for organoid images, with the aim of incorporating into Organotchi those offering the best balance of accuracy, generalizability across imaging conditions, and ease of use. The benchmark included *SegFormer* (Xie *et al*, 2021), *Cellpose* (Stringer *et al*, 2021), *Cellpose-SAM* (Pachitariu *et al*, 2025), *StarDist* (Weigert *et al*, 2020; Schmidt *et al*, 2018), *MOrgAna* (Gritti *et al*, 2021), *OrgLine* (Deng *et al*, 2026), *OrganoID* (Matthews *et al*, 2022), and *μSAM* (Archit *et al*, 2025), and was performed on two brain organoid datasets.

The first dataset, hereafter referred to as testing dataset A, consists of a subset of a publicly available collection of approximately 1,400 images of 64 individually trackable organoids (differentiation according to Lancaster & Knoblich, 2014), imaged at 10 time points over 30 days by two independent laboratories (Lab A, Lab B) (Schröter *et al*, 2024). The dataset provides pixel-level ground-truth masks and the organoids display diverse morphologies and variable image appearance, making it a challenging and realistic test case. Because the plating and imaging conditions often vary depending on the differentiation protocol and the scientific question, we assembled a second dataset, hereafter referred to as testing dataset B (see Methods), comprising ten brightfield images with manually generated ground-truth masks. These images were selected to represent highly distinct imaging scenarios, spanning a range of magnifications, single and multiple organoids per field, and different regional identities, developmental timepoints, and morphologies.

Using cross-validation on the full collection, Schröter and colleagues (Schröter *et al*, 2024) previously showed that *SegFormer* outperformed two classical pipelines, *CellProfiler* and *OrganoSeg* (Stirling *et al*, 2021; Borten *et al*, 2018), as well as the machine-learning tool *MOrgAna* (Gritti *et al*, 2021). Because the model weights were never released, we first reconstructed *SegFormer* using the same MiT-B0 encoder and a similar training regime to that of the original study (Schröter *et al*, 2024) (see Methods). The performance of the reconstructed model, evaluated as mean Intersection over Union (mIoU), mean Dice coefficient (mDice), and pixel-wise accuracy, was consistent with that previously reported (Figure S1). We further added a classical post-processing filter, which corrected segmentation failures caused by imaging artifacts (Figure S2; see Methods).

There are several segmentation tools available, including generalist foundation models, and so we compared *SegFormer*, with and without the post-processing filter, against *Cellpose* (Stringer *et al*, 2021), *Cellpose-SAM* (Pachitariu *et al*, 2025), *StarDist* (Weigert *et al*, 2020; Schmidt *et al*, 2018), *MOrgAna* (Gritti *et al*, 2021), *OrgLine* (Deng *et al*, 2026), *OrganoID* (Matthews *et al*, 2022), and *μSAM* (Archit *et al*, 2025). All tools were applied in a zero-shot setting, with the exception of *MOrgAna*, which does not provide a pretrained model and is instead designed to be trained by the user on a small number of annotated images. MOrgAna was then trained on five images from the original collection that were not included in testing dataset A. For *StarDist*, *Cellpose*, *Cellpose-SAM*, and *μSAM*, whose accuracy is sensitive to expected object-scale parameters, we supplied per-image estimates for each tool, since organoid size changes substantially over the imaged culture period. Each tool was otherwise applied according to its own established workflow (see Methods). As a baseline, we also included a purely classical approach (hereafter *Classical*), based on Otsu intensity thresholding (see Methods).

Segmentation performance was evaluated using the Dice coefficient, the IoU, and the average precision (AP) and F1 score at an IoU threshold of 0.50 (AP50 and F1@50; see Methods).

On testing dataset A, *SegFormer* (both with and without postprocessing filter) and *OrgLine* substantially outperformed every other method (Figure 2A, Table S1). The fourth best performing segmentation tool was *StarDist*, followed by the *Classical* segmentation approach (Figure 2A, Table S1), which was effective as long as organoid and background were sufficiently separated in grayscale intensity, as expected from an approach relying on Otsu thresholding (Figure S3A). The performance of the remaining tools was substantially inferior, with a mDice always lower than 0.800 (Figure 2A and S3A, Table S1).

**Figure 2.**
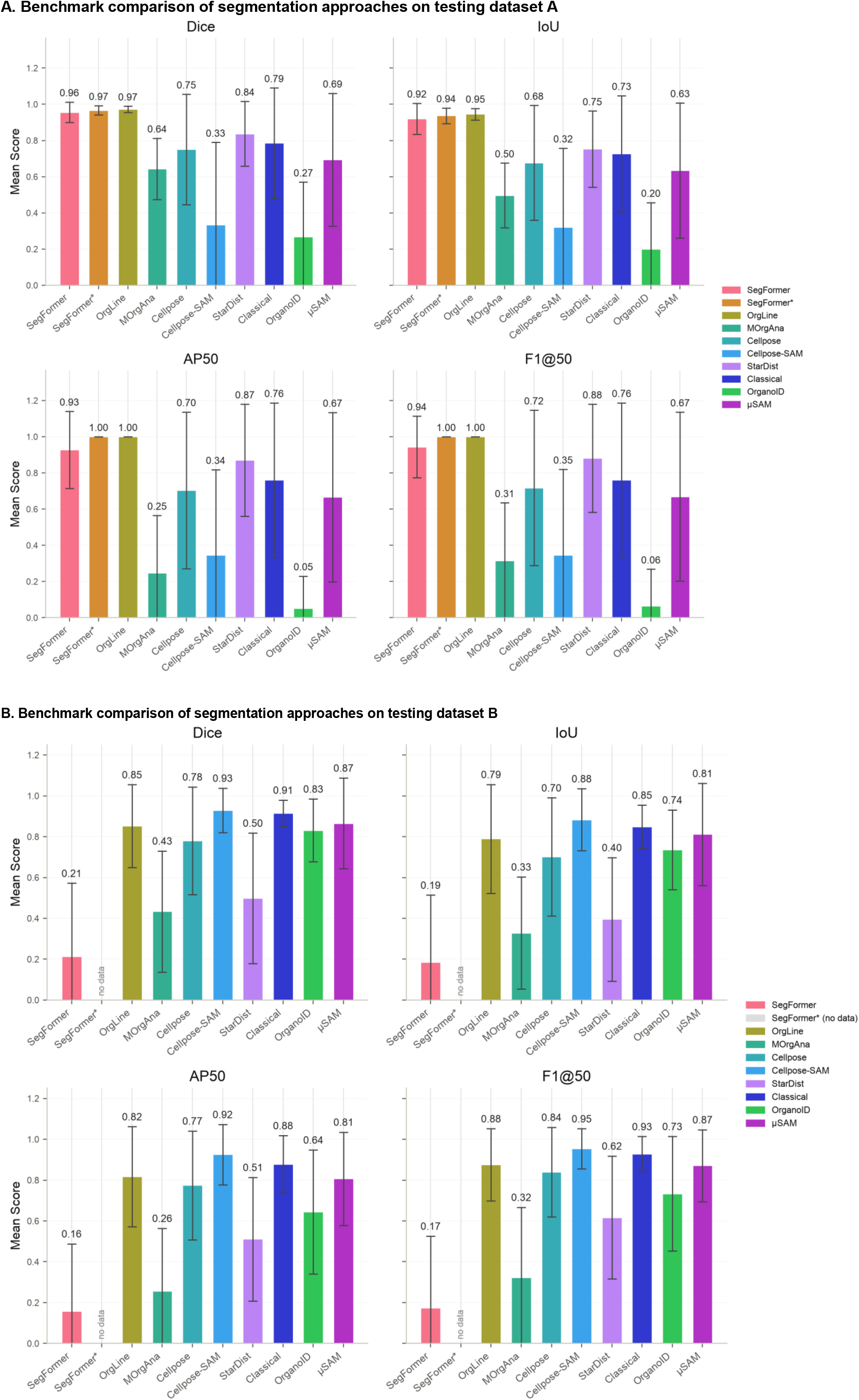
Benchmark comparison of segmentation approaches on (A) testing dataset A and (B) testing dataset B. The panels depict mean ± standard deviation of Dice, IoU, Average Precision at IoU 0.50 (AP50), and F1 score at IoU 0.50 (F1@50), for the *SegFormer*-based model without and with an additional layer of post-processing classical filtering (*SegFormer*\*; only used for testing dataset A), *OrgLine*, *MOrgAna*, *Cellpose*, *Cellpose-SAM*, *StarDist*, *Classical*, *OrganoID*, and *μSAM* (see Methods).

On testing dataset B, *Cellpose-SAM* and *Classical* were the best-performing segmentation methods, both in terms of accuracy and stability (Figure 2B, Table S1). *OrgLine*, *Cellpose*, *OrganoID*, and *μSAM* also performed well, with mDice ranging between 0.780 and 0.865 in all cases (Figure 2B, Table S1). *SegFormer*, *StarDist*, and *MOrgAna* performed poorly in this setting (Figure 2B, Table S1) and segmentation accuracy depended strongly on the individual image (Figure S3B).

Comparing performance across the two datasets showed that no single method performed best in both, and that accuracy on one dataset did not reliably predict accuracy on the other (Figure 2, Table S1). Because Organotchi was intended to perform robustly regardless of the brain organoid protocol and resulting morphology, culture stage, well format, imaging magnification, and number of organoids per field, we prioritized approaches that combined high accuracy with stable performance and required little or no retraining or user input.

On this basis, we selected three approaches (Figure 1): *SegFormer* and *OrgLine* for their near-perfect segmentation on testing dataset A, and the *Classical* approach for its high and stable performance on both datasets, its lack of training requirements, and its near-instantaneous segmentation regardless of image size. Importantly, whereas *SegFormer* performs semantic segmentation and thus merges contiguous organoids, *OrgLine* has the additional advantage of resolving adjacent or overlapping organoids as distinct instances (Deng *et al*, 2026). Offering these complementary routes allows users to choose the one best suited to their data, for example *SegFormer* for individually imaged organoids at 5× magnification, *OrgLine* for fields containing multiple, potentially overlapping organoids, and *Classical* for fast and reliable segmentation across a range of settings.

### Application of Organotchi across experimental contexts

We next tested the generalizability, versatility, and completeness of Organotchi for segmenting and characterizing organoid morphology through four use cases. Throughout this study, we use “morphology” to refer to features measurable from an organoid’s projection in a brightfield image, namely its size and shape.

In all case studies, we applied the analysis pipeline characteristic of Organotchi (Figure 1, see Methods). The segmentation approach used in each case study was chosen in light of the benchmarking results reported above, and is stated explicitly when each case study is introduced. To assess whether morphology alone could distinguish experimental groups, a non-redundant subset of the extracted features (area, perimeter, diameter, circularity, roundness, skeleton length, aspect ratio, maximum Feret diameter, solidity, and eccentricity; see Methods) was embedded in two dimensions using UMAP, and organoids were clustered with Hierarchical Density-Based Spatial Clustering of Applications with Noise (HDBSCAN). HDBSCAN was selected for not requiring a predefined number of clusters, for accommodating clusters of varying shape and density, and for classifying outlying organoids as noise (Campello *et al*, 2013). Cluster assignments were then compared with the experimental grouping variable using four complementary measures. The adjusted Rand index (ARI) and normalized mutual information (NMI) quantify overall agreement between clusters and groups. Both equal 1 for a perfect match, while values close to 0 indicate agreement no better than random (ARI) or no shared information (NMI). Weighted cluster purity measures how strongly each cluster is dominated by a single group. Finally, the chi-square test of independence assesses whether the association between cluster assignment and group label is statistically significant. Outliers were flagged by Organotchi using Tukey’s interquartile-range (IQR) rule, applied independently within each group, imaging day, and morphological feature, and excluded from downstream analysis. The percentage of outliers was quantified for each case study and reported in the supplemental tables (Tables S2–S5).

### Case study 1: characterizing growth and morphology in patient-derived neural organoids

The dataset used in the context of the training of *SegFormer* encompasses images of neural organoids differentiated from induced pluripotent stem cells (iPSCs) generated from peripheral blood mononuclear cells of a healthy control (wt2D), two patients with TUBA1A-(A1A-1) and TUBB2A-associated tubulinopathy (B2A-2), and one patient with the neurotransmitter disorder tyrosine hydroxylase (TH) deficiency (TH2-7) (Schröter *et al*, 2024). Sixteen organoids per clone per lab were tracked across days 2-30, and imaged independently by two laboratories (Lab A, Lab B). Differentiation took place in 96-well plates, using an unguided protocol (Lancaster & Knoblich, 2014), and imaging was performed at 5× magnification. In the original study, this dataset served primarily as a benchmark for segmentation methods, and morphological analysis was limited to organoid size over time and a principal component analysis of clone diversity (Schröter *et al*, 2024). We therefore used it as a first test case for Organotchi, extending this characterization to a broader set of morphological features and their rates of change, and statistical comparisons between clones. Within Organotchi, we used *SegFormer* to obtain organoid masks (Figure 3A and S4), as this dataset corresponds to its training domain.

**Figure 3.**
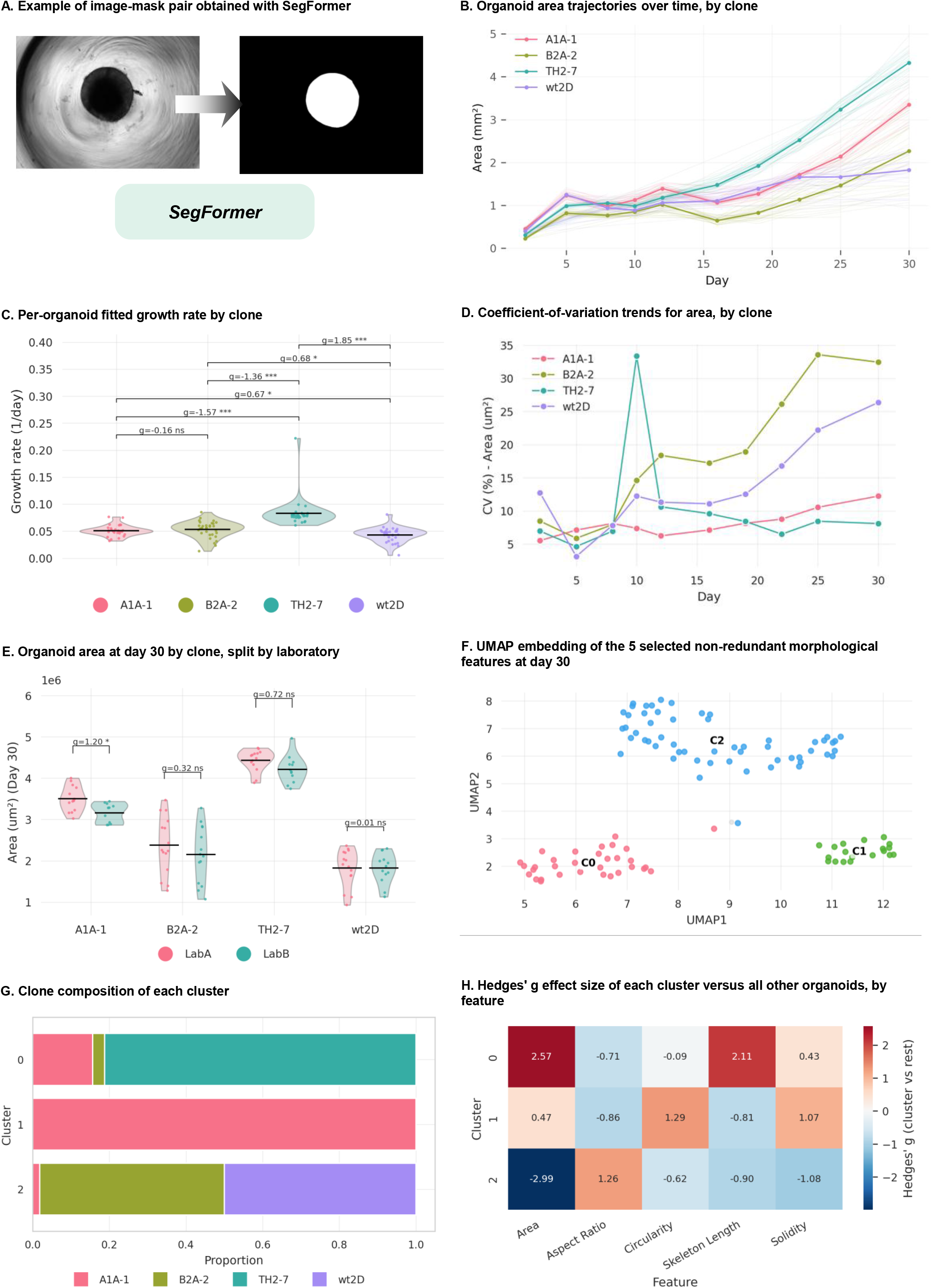
Case study 1: brain organoid morphology across four clones (wt2D = healthy control; A1A-1 = TUBA1A tubulinopathy; B2A-2 = TUBB2A tubulinopathy; TH2-7 = tyrosine-hydroxylase deficiency), tracked across days 2-30, imaged independently by two laboratories (Lab A, Lab B). (A) Example of image-mask pair obtained with *SegFormer*. (B) Individual-organoid area trajectories over time (thin lines), by clone, with the clone-level mean overlaid (thick line). (C) Per-organoid fitted growth rate k (exponential model, Area(t) = Area₀×exp(k×t)), by clone, with pairwise statistics. (D) Coefficient-of-variation trends for area, by clone. (E) Area at day 30 by clone, split by laboratory (Lab A vs. Lab B), with pairwise statistics restricted to the Lab A-Lab B comparison within each clone. (F) UMAP embedding of the 5 selected non-redundant morphological features at day 30 (n=108 organoids), colored by HDBSCAN cluster (C0-C2; grey = unassigned/noise). (G) Clone composition of each cluster. (H) Hedges’ g effect size of each cluster versus all other organoids, by feature. Pairwise comparisons throughout use two-sided Welch’s t-tests with Hedges’ g effect sizes, Benjamini-Hochberg FDR-corrected across all comparisons within each panel: * FDR < 0.05, ** FDR < 0.01, *** FDR < 0.001, ns = not significant (FDR ≥ 0.05).

Analysis of organoid growth by clone (Figure 3B, 3C, Table S2A) showed TH2-7 reaching the highest mean area by day 30, followed by A1A-1, B2A-2, and wt2D. Fitted per-well growth rates averaged 0.083/day for TH2-7, 0.053/day for B2A-2, 0.051/day for A1A-1, and 0.043/day for wt2D (Figure 3C). Variability in area also differed markedly between clones. The coefficient of variation rose almost monotonically over time in B2A-2 and wt2D, peaking at day 30 in B2A-2, whereas it remained low throughout in TH2-7 and A1A-1 (Figure 3D). Final mean organoid area agreed closely between laboratories within every clone except A1A-1, for which organoids were slightly larger in Lab A (Figure 3E). The specific ranking of clones and their size differences were nonetheless preserved in both laboratories (Figure 3B, 3E).

HDBSCAN clustering identified three clusters from the four clone categories (weighted purity 0.682; Χ^2^=168.0, P-value<10^−4^; ARI=0.555 and NMI=0.679; n=108 organoids, 107 assigned to a cluster and 1 classified as noise; Figure 3F), indicating that clone identity is only partially recoverable from morphology alone. Cluster 1 contained only A1A-1 organoids and had the highest circularity and solidity (Figure 3G, 3H, Table S2B). Cluster 0 was predominantly TH2-7 and was characterized by larger area and skeleton length (Figure 3G, 3H, Table S2B). Cluster 2, the largest cluster, was a near-even mix of wt2D and B2A-2 organoids, with a small A1A-1 contribution, and was distinguished by smaller area and larger aspect ratio (Figure 3G-H, Table S2B).

Taken together, the growth trajectories reported here match those originally described for this dataset (Schröter *et al*, 2024) (Figure 3B). Beyond replicating these growth-based findings, however, the expanded feature panel and unsupervised clustering resolve the two tubulinopathy clones as morphologically distinct from one another (Figure 3G, 3H). Specifically, while A1A-1 organoids formed a clone-pure cluster defined by increased circularity and solidity, the only cluster with B2A-2 organoids carried the opposite signature (Figure 3G, 3H). B2A-2 organoids were also substantially smaller by day 30 (Figure 3B), showed greater within-clone variability in size than any other group (Figure 3D), and were morphologically more similar to controls than any other disease clone (Figure 3G).

### Case study 2: pharmacological modulation of Wnt signaling in dorsal forebrain organoids

Regional identity in neural organoids is established by the same morphogen gradients that pattern the neural tube in vivo, and guided protocols exploit this by supplying patterning cues exogenously (Kelley & Pașca, 2022; Eichmüller & Knoblich, 2022). Wnt signaling is central to this process, specifying position along the anterior-posterior axis, with dorsal forebrain identity depending on Wnt inhibition at the anterior pole (Qian *et al*, 2016; Paşca *et al*, 2015). Beyond patterning, canonical Wnt signaling regulates neural progenitor proliferation and the timing of neurogenesis, so its manipulation is expected to affect not only regional fate but also tissue growth (Hirabayashi *et al*, 2004; Chenn & Walsh, 2002).

To examine Wnt-dependent effects on early forebrain organoid growth and general morphology, we applied a simple pharmacological perturbation. Forebrain organoids (Velasco *et al*, 2019) were cultured in 96-well plates under three conditions: CHIR99021 (CHIR), a GSK3 inhibitor widely used to activate canonical Wnt signaling; ICG-001 (ICG-1), an inhibitor of Wnt/β-catenin/TCF-mediated transcription; and DMSO vehicle as control. We expected canonical Wnt activation to increase organoid growth through enhanced progenitor proliferation, and Wnt inhibition to reduce it relative to control (Hirabayashi *et al*, 2004; Chenn & Walsh, 2002). Imaging was performed from day 2 to day 12, using a 4× magnification. As in the preceding case study, masks were obtained with *SegFormer*, as the imaging conditions, with individually cultured forebrain organoids imaged at 4× magnification, closely matched those of its training domain (Figure 4A and S5).

**Figure 4.**
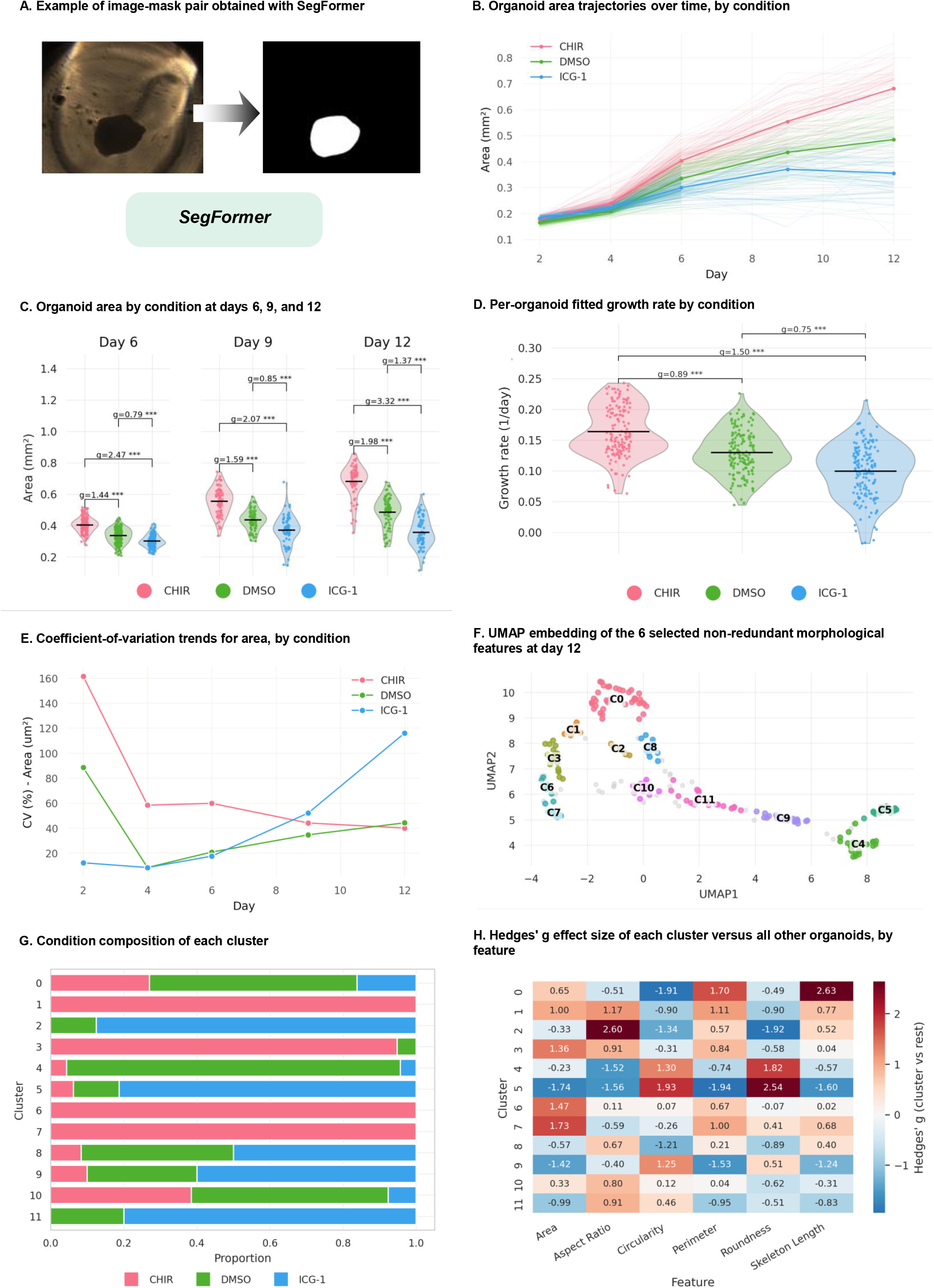
Case study 2: pharmacological modulation of Wnt signaling in dorsal forebrain organoids (CHIR: CHIR99021 = GSK3 inhibitor/Wnt activator; DMSO = vehicle control; ICG-1: ICG-001 = beta-catenin/CBP inhibitor), tracked across days 2-12. (A) Example of image-mask pair obtained with *SegFormer*. (B) Individual-organoid area trajectories over time (thin lines), by condition, with the condition-level mean overlaid (thick line). (C) Area by condition at days 6, 9, and 12, with pairwise statistics. (D) Per-organoid fitted growth rate k (exponential model, Area(t) = Area₀×exp(k×t)), by condition, with pairwise statistics. (E) Coefficient-of-variation trends for area, by condition. (F) UMAP embedding of the 6 selected non-redundant morphological features at day 12 (n=224 organoids), colored by HDBSCAN cluster (C0-C11; grey = unassigned/noise). (G) Condition composition of each cluster. (H) Hedges’ g effect size of each cluster versus all other organoids, by feature. Pairwise comparisons throughout use two-sided Welch’s t-tests with Hedges’ g effect sizes, Benjamini-Hochberg FDR-corrected across all comparisons within each panel: * FDR < 0.05, ** FDR < 0.01, *** FDR < 0.001, ns = not significant (FDR ≥ 0.05).

Organoid growth trajectories separated early on, with divergence between conditions increasing over time (Figure 4B, 4C, Table S3A). CHIR-treated organoids reached the highest mean area of the three conditions at day 12, followed by DMSO and ICG-1-treated organoids; notably, the ICG-1 mean growth trajectory plateaued and slightly declined between day 9 and day 12 (Figure 4B, 4C). Per-well fitted growth rates also differed significantly between conditions and were particularly variable for ICG-1-treated organoids, with some organoids showing near-zero growth (Figure 4D). Consistent with this, the coefficient of variation (CV) trends for area rose through day 12 for ICG-1-treated organoids while remaining comparatively low from day 2 onwards for the two other conditions (Figure 4E).

HDBSCAN clustering at day 12 identified 12 clusters that only partially recovered the three-way condition label (ARI=0.142, NMI=0.301), even though a chi-square test rejected independence between cluster and condition (weighted purity 0.756, Χ^2^=191.6, P-value<10^−4^; n=224 organoids, 193 assigned to a cluster, 31 classed as noise; Figure 4F). At one extreme, clusters 1, 3, 6, and 7 were composed entirely of CHIR-treated organoids, while clusters 2 and 11 had none, and clusters 4, 5, and 8 had a single CHIR-treated organoid each (Figure 4G). Larger area and perimeter were the features that most consistently separated the CHIR-from the ICG-1/DMSO-enriched clusters and the remaining morphological features were not clearly condition-specific, as they often characterized clusters of mixed composition (Figure 4G, 4H, Table S3B).

Together, these results indicate that pharmacological modulation of canonical Wnt signaling produces a growth-dominant morphological phenotype in early forebrain organoids. Specifically, CHIR-treated organoids grew fastest and reached the largest size, ICG-1-treated organoids grew slowest and became progressively more heterogeneous, and DMSO controls fell consistently in between. This is consistent with the established role of canonical Wnt signaling in setting cortical progenitor output (Hirabayashi *et al*, 2004; Chenn & Walsh, 2002), and suggests that, at least during the considered developmental window, the influence of Wnt on tissue growth may be readily and robustly captured by morphometric profiling. Determining whether Wnt modulation additionally influenced the regional identity of the organoids, in line with its role in neural patterning (Qian *et al*, 2016; Paşca *et al*, 2015), would require complementary approaches, such as immunostaining for regional markers or RNA sequencing.

### Case study 3: response to 5-FU in MCF-7 spheroids

The segmentation approaches incorporated into Organotchi were selected and benchmarked exclusively on brightfield images of neural organoids. To test whether Organotchi generalizes beyond this development domain, we used an imaging dataset of MCF-7 breast cancer spheroids from a recently published study (Raic *et al*, 2026). These spheroids differ from neural organoids in origin and cellular composition, forming by overnight self-aggregation of an immortalized luminal A breast cancer line rather than by stem cell differentiation, and in size, compactness, and optical appearance (Figure S6).

Raic and colleagues combined time-resolved ¹H NMR exometabolomics with morphological, transcriptional, and viability readouts, comparing 2D monolayers and 3D spheroids exposed to 5-fluorouracil (5-FU), a widely used chemotherapy medication, at approximately its 2D IC₅₀ (220 µM) over 72 h (Raic *et al*, 2026). Morphometric measurements were obtained by manual annotation in ImageJ/FIJI, yielding three parameters for at least 12 spheroids per condition and time point across three independent experiments. This dataset therefore allows a direct test of whether Organotchi recovers morphometric results obtained manually. It also allows the reported morphology-metabolism relationships to be revisited with a broader panel of morphological features. Brightfield images were acquired at 0, 24, 48, and 72 h post-treatment with a 4× objective in 96-well plates. This plating strategy and magnification resembled that of the training data of *SegFormer*, but the appearance of the spheroids was markedly different. These images were therefore segmented using the *Classical* segmentation approach, which performed reliably in our benchmark when objects showed strong intensity contrast against the background (Figure 2 and S3), which was the case here (Figure 5A and S6).

**Figure 5.**
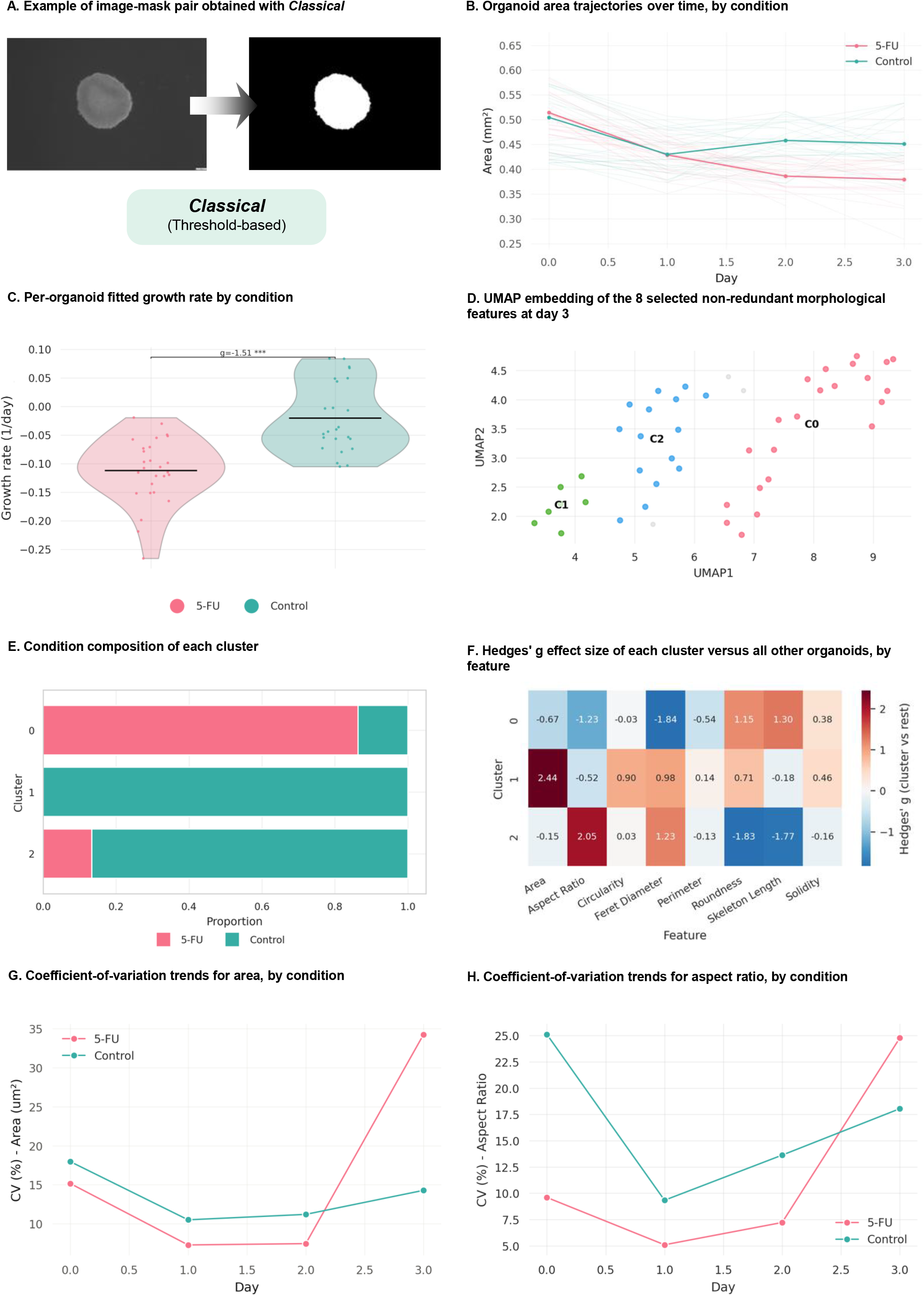
Case study 3: exposure response of MCF-7 spheroids to 5-FU vs. Control, tracked across days 0-3. (A) Example of image-mask pair obtained with *Classical*. (B) Individual-organoid area trajectories over time (thin lines), by condition, with the condition-level mean overlaid (thick line). (C) Per-organoid fitted growth rate k (exponential model, Area(t) = Area₀×exp(k×t)), by condition, with pairwise statistics. (D) UMAP embedding of the 8 selected non-redundant morphological features at day 3 (n=47 organoids), colored by HDBSCAN cluster (C0-C2; grey = unassigned/noise). (E) Condition composition of each cluster. (F) Hedges’ g effect size of each cluster versus all other organoids, by feature. (G) Coefficient-of-variation trends for area and (H) aspect ratio, by experimental condition. Pairwise comparisons throughout use two-sided Welch’s t-tests with Hedges’ g effect sizes, Benjamini-Hochberg FDR-corrected across all comparisons within each panel: * FDR < 0.05, ** FDR < 0.01, *** FDR < 0.001, ns = not significant (FDR ≥ 0.05).

The mean area of the spheroids decreased from day 0 to day 1 in both conditions, diverging from day 1 onwards (Figure 5B). Mean area of control (non-treated) spheroids increased slightly by day 2 and remained stable through day 3, whereas that of 5-FU-treated spheroids continued declining throughout (Figure 5B, Table S4A). These differences were non-significant at day 1 but became significant at days 2 and 3 (Table S4A). Per-well fitted growth rates were also significantly different between conditions (Figure 5C). Notably, every single 5-FU-treated spheroid had a negative growth rate, compared to 7 of 24 Control spheroids that still showed positive growth (Figure 5C). Organoid diameter and maximum Feret diameter followed a trajectory very similar to area. Differences in aspect ratio, skeleton length, and eccentricity became significant only at day 3, while the remaining shape descriptors (circularity, roundness) and solidity never reached significance at any time point (Table S4A).

HDBSCAN clustering at day 3 identified three clusters showing moderate correspondence with the Control/5-FU label (ARI=0.422, NMI=0.423), while a chi-square test rejected cluster-condition independence decisively (weighted purity 0.886, Χ^2^=26.7, P-value<10^−4^; n=47 spheroids, 44 assigned to a cluster and 3 classified as noise; Figure 5D). Cluster 1 was condition-pure, composed solely of Control spheroids and characterized by markedly larger area (Figure 5E-F, Table S4B). Clusters 2 and 0 were, on the other hand, composed of a mix of spheroids from the two conditions, albeit in very distinct proportions (Figure 5E, 5F, Table S4B). Cluster 2, which was mostly composed of Control spheroids and characterized by larger aspect ratio (Figure 5E, 5F, Table S4B). Cluster 0, mostly composed of 5-FU-treated spheroids, had the opposite signature and was further characterized by increased roundness and larger skeleton length (Figure 5E, 5F, Table S4B).

When examining the CV of the two morphological parameters that most clearly distinguished Control spheroids by day 3, namely area and aspect ratio, we found that both were higher in Control than 5-FU at every time point but day 3, when they increased considerably in 5-FU-treated spheroids relative to the previous days (Figure 5G, 5H).

Taken together, this case study shows that Organotchi transfers to a 3D culture system well outside its development domain and recovers the morphometric conclusions of the original study (Raic *et al*, 2026). Specifically, untreated MCF-7 spheroids decreased slightly in size before re-expanding and plateauing, whereas 5-FU-treated spheroids underwent a progressive reduction of roughly 25% in projected area and 12% in Feret diameter over 72 h. Importantly, however, the rising CV of area and aspect ratio in the 5-FU group at day 3 adds a dimension not captured by the original study. Treated spheroids were the more homogeneous group for most of the time points, yet became markedly more variable in size by the final time point, with some shrinking substantially while others remained comparable to controls. This suggests that, despite the consistent group-level size and shape effects reported, the magnitude of the 5-FU response varies considerably between individual spheroids. This is a form of heterogeneity that group-mean comparisons cannot reveal, and that may be informative as a readout of drug efficacy.

### Case study 4: cystic morphology in choroid plexus organoids seeded in two media conditions

The choroid plexus is a secretory epithelium of the brain ventricles that produces cerebrospinal fluid (CSF) and forms the blood-CSF barrier (Fame & Lehtinen, 2020; Lehtinen *et al*, 2013; Ghersi-Egea *et al*, 2018). Choroid plexus organoids recapitulate this identity in vitro, forming a polarized epithelium enclosing fluid-filled compartments (Pellegrini *et al*, 2020). Cyst-like cavities are therefore an expected and desired feature of these cultures, in contrast to cerebral organoid protocols, where their appearance signals off-target differentiation (Ikeda *et al*, 2024). Thus, the same morphological feature carries opposite interpretations depending on protocol, which makes cystic area a useful test of whether an interpretable feature panel supports protocol-specific quality criteria.

This case study compares choroid plexus organoids (Pellegrini *et al*, 2020) derived under two media conditions: a batch of iPSCs was seeded in iPSC media (mTesR Plus) and another directly in the protocol’s original differentiation media. Organoids were imaged in two regimes: individually in 96-well plates from day 2 to day 7 using a 4× magnification, and, after replating at day 10, in 6-well plates from day 13 to day 30, acquired as 5×5 stitched tile grids with a 1.25× magnification. In both phases, brightfield images were segmented with *OrgLine,* given its good performance on both benchmarking datasets (Figure 2 and S3) and its particular advantage of resolving adjacent organoids as separate instances (Figure 6A and S7).

**Figure 6.**
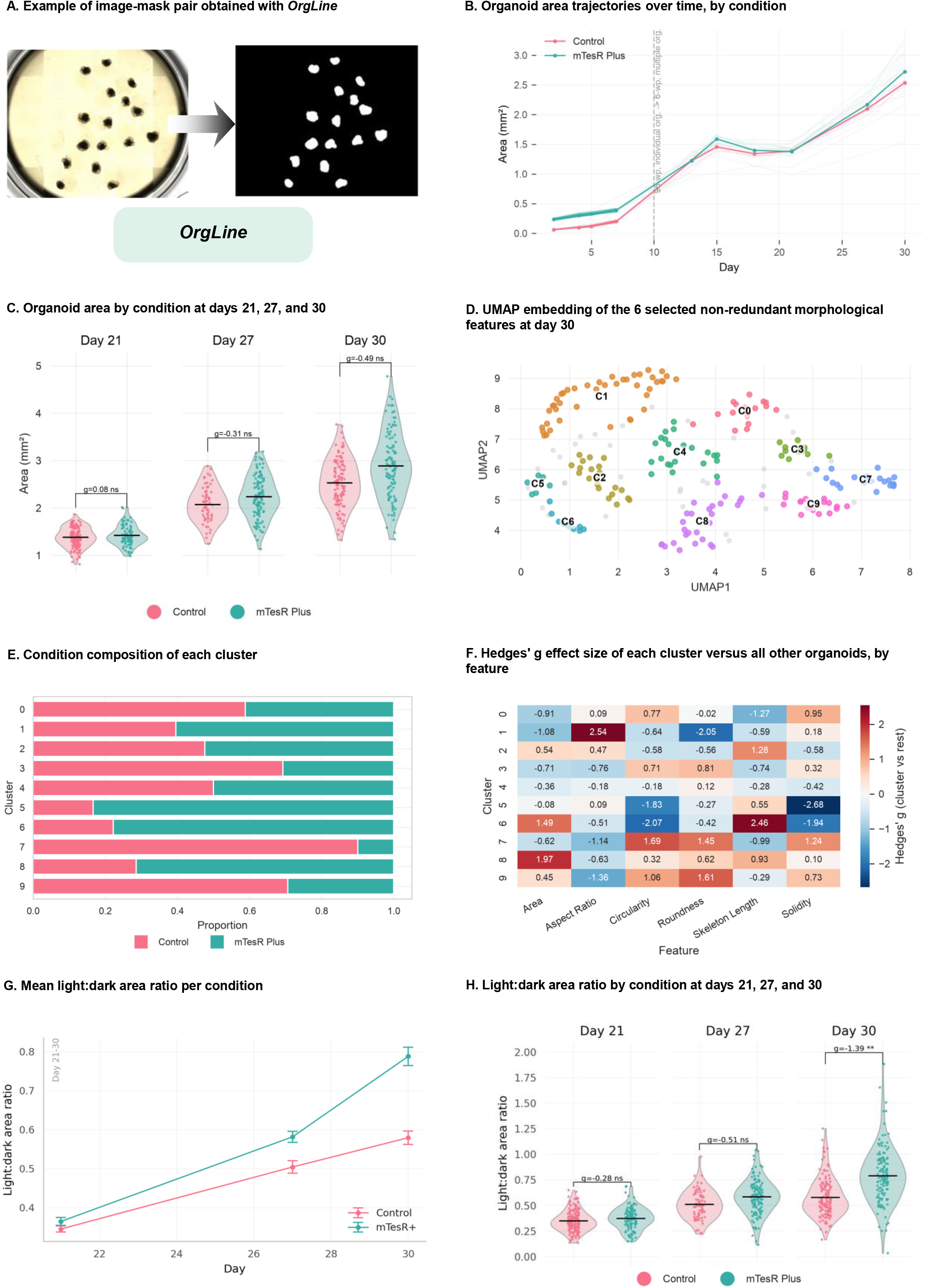
Case study 4: Effects of seeding media in choroid-plexus organoid differentiation (mTesR Plus vs. Control), tracked across days 2-30. (A) Example of image-mask pair obtained with *OrgLine*. (B) Individual-organoid area trajectories over time (thin lines), by condition, with the condition-level mean overlaid (96-well-to-6-well plate transition marked at day 10). (C) Area by condition at days 21, 27, and 30, with pairwise testing. (D) UMAP embedding of the 6 selected non-redundant morphological features at day 30 (n=231 organoids), colored by HDBSCAN cluster (C0-C9; grey = unassigned/noise). (E) Condition composition of each cluster. (F) Hedges’ g effect size of each cluster versus all other organoids, by feature. (G) Mean light:dark area ratio per condition over the Day 21-30 developmental window (mean ± SEM). (H) Light:dark area ratio by condition at days 21, 27, and 30 (Day 21-30 window), with pairwise statistics. Pairwise comparisons throughout use two-sided Welch’s t-tests with Hedges’ g effect sizes, Benjamini-Hochberg FDR-corrected across all comparisons within each panel: * FDR < 0.05, ** FDR < 0.01, *** FDR < 0.001, ns = not significant (FDR ≥ 0.05).

Growth profiles did not differ substantially between conditions by the end of the differentiation period (Figure 6B, C, Table S5A), but within-condition area variability increased markedly throughout, especially in mTesR Plus organoids (Figure 6C). HDBSCAN clustering at day 30 identified ten clusters (n=231 organoids, 198 assigned to clusters and 33 classified as noise; Figure 6D) and correspondence with condition was extremely weak by ARI/NMI (ARI=0.022, NMI=0.059; weighted purity 0.657; χ²=30.6, P-value=3.5×10⁻⁴). There were no condition-pure clusters, and only a subset of organoids from each condition was clearly distinguishable from the rest (Figure 6E). Clusters 3, 7, and 9 were predominantly Control and were all characterized by increased circularity and roundness (Figure 6E, 6F, Table S5B). At the other end of the spectrum, clusters 5, 6, and 8 were predominantly mTesR Plus, with 5 and 6 showing reduced circularity and solidity, and 6 and 8 being substantially larger and having increased skeleton length (Figure 6E, 6F, Table S5B). The remaining four clusters (0, 1, 2, 4) accounted for approximately 53.5% of organoids and were composed of an approximately even mix of Control and mTesR Plus organoids, driving the weak ARI/NMI despite the significant overall association (Figure 6E). We thus conclude that the differences between conditions in organoid morphology (Table S5A) are mostly driven by a subset of organoids and do not represent a general, condition-specific trend.

Because fluid-filled cysts are a defining feature of choroid plexus organoids, we next used Organotchi to track them and quantify their area throughout the culture period. For this purpose, Organotchi offers an optional analysis that complements the standard feature set by partitioning each organoid into lighter and darker regions by Otsu thresholding (Figure S8, see Methods). In brightfield, fluid-filled cysts transmit more light than surrounding tissue, so the light fraction provides a proxy for cystic area and the dark fraction for dense cellular tissue. We report their ratio as the light:dark ratio.

Cysts were apparent on visual inspection at day 21, and we therefore compared the light:dark ratio at days 21, 27, and 30. Both conditions began from near-identical values and increased over the window, diverging significantly by day 30 (Figure 6G, 6H). The rate of change was also different across conditions, with the increase in Control decelerating over the final interval (+46% from day 21 to 27, +15% from day 27 to 30), whereas mTesR Plus continued to rise (Figure 6G).

Organotchi, while allowing for the longitudinal tracking of individual organoids, also makes it possible to track organoid count per well over the developmental timeline. At day 13, the first day of imaging after organoid replating, segmentation with *OrgLine* resulted in only 2.5 organoids/well on average for Control versus 6.8/well for mTesR Plus; by day 15 the two conditions converged (10.2 vs. 9.9/well). A second, larger divergence appears at day 27, where the Control count drops sharply to 6.3/well while mTesR Plus rises to 13.0/well, before both converge again by day 30 (11.7 vs. 12.0/well). This suggests that at days 13 and 27, a considerable proportion of organoids may have gone undetected by *OrgLine*.

As previously discussed, the segmentation results obtained from different tools largely depended on imaging and organoid-culturing settings. We therefore asked whether the results of this case study, particularly organoid counts, depend on the segmentation method used. We repeated the analysis using *SegFormer* for the early phase (days 2-7) and the *Classical* segmentation approach from day 13 onward.

During the single-organoid phase (days 2-7), *OrgLine* and *SegFormer* detected equivalent numbers of organoids: always exactly one per well (Figure 7A). In the multi-organoid phase, however, the *Classical* approach generally detected more objects than *OrgLine*, reaching statistical significance at days 13 and 15, while *OrgLine* detected significantly more organoids than the *Classical* approach only at day 30 (Figure 7A). Interestingly, the difference between conditions in detected organoid counts was also smaller with the *Classical* approach than with *OrgLine* (Figure 7B). Without ground-truth annotations for these images, we cannot determine which segmentation method is more accurate overall, since a higher count may reflect improved detection, fragmentation of individual organoids into multiple objects, or misdetection of imaging artifacts (such as agglomerates of dead cells or air bubbles) as organoids. However, direct visual inspection of the five images with the lowest *OrgLine*-detected counts from day 13 onwards (ties broken at random with a fixed seed) confirmed clear under-detection by *OrgLine* and generally robust segmentation with the *Classical* approach (Figure S9). In every one of the five cases, *OrgLine* detected at most one organoid (including one image where no objects were detected), while the *Classical* approach identified 8-14 candidate organoids in the identical raw image (Figure S9). In some cases, image artifacts were indeed segmented as organoids by the *Classical* approach, but these were correctly flagged as outliers owing to their substantially distinct morphology, so they did not compromise the quality or robustness of the analysis.

**Figure 7.**
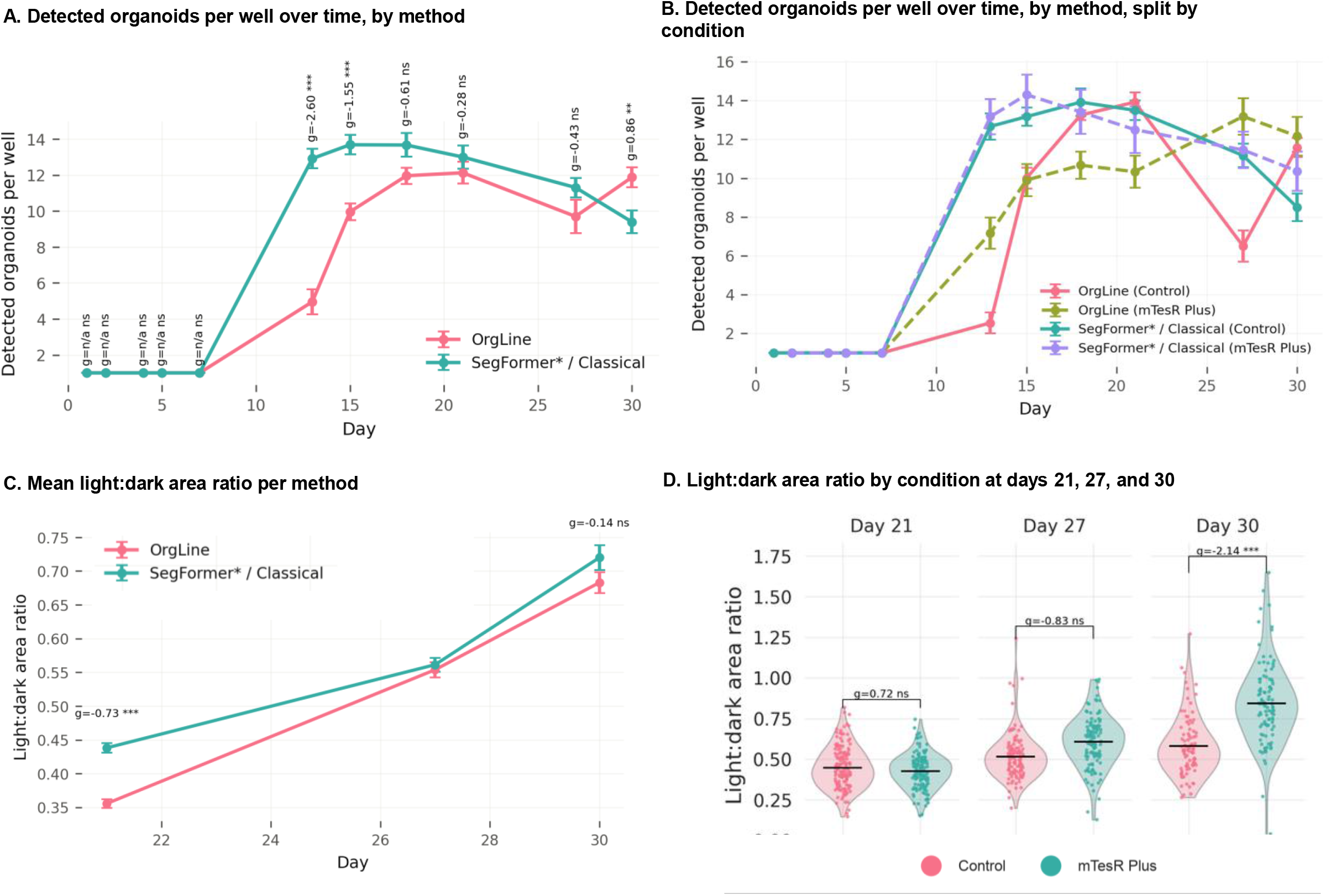
Segmentation-method robustness check for case study 4 *OrgLine* vs. the alternative pipeline consisting of *SegFormer*\* (*SegFormer* with postprocessing segmentation filter) for the single-organoid phase (days 2-7) and the *Classical* segmentation approach from day 13 onward. (A) Detected organoids per well over time, by method (both conditions pooled), with pairwise statistics per day. (B) The same counts, split by method and condition (C) Mean light:dark area ratio per method over the Day 21-30 developmental window (mean ± SEM; both conditions pooled), with pairwise statistics per day. (D) Light:dark area ratio by condition at days 21, 27, and 30 (Day 21-30 window), with pairwise statistics, as assessed with the *Classical* segmentation approach. Pairwise comparisons throughout use two-sided Welch’s t-tests with Hedges’ g effect sizes, Benjamini-Hochberg FDR-corrected across all comparisons within each panel: * FDR < 0.05, ** FDR < 0.01, *** FDR < 0.001, ns = not significant (FDR ≥ 0.05).

Concerning the light:dark ratio, the *Classical* segmentation detected a larger “light” organoid area than *OrgLine*’s at day 21, but the difference between the two methods from day 27 onward, was non-significant (Figure 7C). Notably, the same pattern in light:dark area ratio between the two conditions that we had previously reported (Figure 6H) was still observed with the *Classical* segmentation approach: mTesR Plus organoids had a statistically significant higher ratio by day 30, but not at days 21 or 27 (Figure 7D).

Taken together, the medium in which the iPSCs were seeded at the start of organoid differentiation had little effect on the overall size and shape of the resulting choroid plexus organoids. However, organoids derived from iPSCs seeded in mTeSR Plus showed more variable morphology and a significantly larger light-to-dark area ratio, indicating a greater cystic fraction, and this difference was consistent across two distinct segmentation methods. This indicated that culturing iPSCs in mTeSR Plus may favor cyst formation in choroid plexus organoids.

## Discussion

### Our contribution: bridging segmentation, morphological characterization, and quality control in organoid research

Neural organoids are increasingly used to model tissue development and disease, and organoid-based models are expected to inform preclinical and regulatory decisions (European Commission, 2026; U.S. Food and Drug Administration (FDA), 2025). Their value for these applications, however, depends on how consistently they can be generated and assessed. Guided protocols can now generate neural organoids with reproducible cell-type composition (Velasco *et al*, 2019; Yoon *et al*, 2019), yet variability between organoids, differentiation batches, and laboratories remains a central concern (Jensen & Little, 2023). This variability arises both from differences between stem cell lines (Boerstler *et al*, 2025; Branco *et al*, 2026) and from non-genetic sources such as differentiation batch, the state of the starting stem cells, and technical handling (Glass *et al*, 2024). In response, consensus perspectives and society standards have called for rigorous experimental design, quantitative quality control, and transparent reporting (Ludwig *et al*, 2023; Sandoval *et al*, 2024; Pașca *et al*, 2025).

Meeting these calls in routine practice requires tools that make quality assessment quantitative, fast, and accessible to laboratories without dedicated image-analysis expertise. This need is becoming more pressing as organoid differentiation is scaled up and automated, with microcavity arrays, liquid-handling robotics, and microfluidic platforms now generating hundreds to thousands of organoids per experiment for screening and disease modelling (Ramani *et al*, 2024; Schuster *et al*, 2020; Brandenberg *et al*, 2020; Renner *et al*, 2020). At this scale, manual inspection becomes a bottleneck. Brightfield imaging offers a practical basis for automated assessment, because it is non-invasive, label-free, and can be repeated on the same organoids throughout the culture period. Yet, criteria for selecting organoids for analysis often remain undefined, although morphological parameters such as the Feret diameter have recently been proposed as indicators of brain organoid quality (Boerstler *et al*, 2025). Meanwhile, existing brightfield image analysis tools focus primarily on segmentation and basic morphology assessment (Borten *et al*, 2018; Kassis *et al*, 2019; Gritti *et al*, 2021; Lefferts *et al*, 2024), and those that track individual organoids over time (Matthews *et al*, 2022) do not assess variability within and between batches or support quality-control decisions.

We developed Organotchi, an open-source tool for the automated segmentation, morphological characterization, and longitudinal tracking of organoids (Figure 1) to address this gap. Organotchi complements three high-performing segmentation routes with morphological characterization and experiment-level quality control. It reports per-feature coefficients of variation by time point and condition and flags outliers that users can exclude from subsequent morphometric analyses. Because organoid identity is tracked throughout, this outlier detection also provides quantitative grounds for excluding specific organoids from continued culture and follow up experiments, and for selecting those that are morphologically representative of their batch. Together, these features make the selection and exclusion of organoids more objective and transparent, in line with recommendations for transparent reporting in preclinical and stem cell research (Landis *et al*, 2012; Ludwig *et al*, 2023). It further tests whether morphology alone distinguishes experimental groups by comparing HDBSCAN clusters with group labels using the adjusted Rand index, normalized mutual information, weighted cluster purity, and a chi-square test. Additionally, to complement the analyses on organoid size and shape, Organotchi can also partition each organoid into lighter and darker regions, an approach we validated for separating cystic areas from dense tissue in choroid plexus organoids 11/5*/2026* 6:31:00 AM, as shown in case study 4. To our knowledge, these components have not previously been combined in a single open-source tool validated on neural organoids across diverse imaging conditions and experimental settings.

Alongside the tool, we release two previously unpublished brightfield datasets of dorsal forebrain and choroid plexus organoids, comprising more than 3,500 raw images and the corresponding binary masks. We also release the trained *SegFormer* checkpoints, which can serve as a starting point for fine-tuning the model to different organoid types and imaging conditions.

### Selection of Organotchi’s segmentation routes through near-zero-shot benchmarking

Organotchi offers three distinct segmentation routes – *OrgLine*, *SegFormer*, and *Classical* –, which were chosen after benchmarking several segmentation methods on two markedly different brightfield datasets of neural organoids under a near-zero-shot setting, that is, without fine-tuning when possible and with at most minimal task-specific input. Because many wet labs lack image-mask pairs of their own organoids, we designed the comparison to reflect a realistic scenario in which each tool received minimal input but no resource-intensive, time-consuming training. We therefore acknowledge that the performance reported here does not necessarily reflect the best possible performance of each tool, and that, for some tools, results could likely improve with fine-tuning.

To account for the specific requirements of each tool, we trained *MOrgAna* on five brain organoid images, while for *StarDist*, *Cellpose*, *Cellpose-SAM*, and *μSAM*, we provided per-image estimates of object size. Without fine-tuning or user input, *OrgLine* and *SegFormer* (with postprocessing filter) achieved mDice scores of 0.97 on the first dataset (Figure 2A, Table S1). On the second, the *Classical* approach, which has minimal input requirements and short running times, performed close to the best-performing tool, *Cellpose-SAM* (Figure 2B, Table S1). Although Cellpose-SAM achieved a slightly higher mDice score (by 1.4 percentage points, with a larger standard deviation), its running time does not scale as well as that of the *Classical* approach, and it required an estimate of object size. These two factors are particularly limiting for long-running, large-scale experiments where hundreds to thousands of brightfield microscopy images are generated and where organoid size changes considerably over time.

### Limitations of the study

Organotchi was developed and benchmarked on brightfield images of neural organoids, and its application outside this domain has so far been tested on a single dataset of MCF-7 breast cancer spheroids. Therefore, its performance on other 3D culture systems remains to be tested. Several features of the current implementation, however, are expected to support such extension. On the one hand, the *Classical* approach combines fixed or Otsu thresholding, applicable to both brightfield and darkfield images, with user-customizable size and circularity filters. On the other hand, *OrgLine*, was originally validated on intestinal, colon, and pancreatic ductal adenocarcinoma datasets alongside the brain dataset from Schröter and colleagues (Schröter *et al*, 2024; Deng *et al*, 2026). Additionally, users can also supply their own SegFormer checkpoint trained on a different image domain, after which Organotchi’s morphological analysis pipeline can be applied without modification.

Similarly, the light/dark partitioning of segmented objects was validated only for separating cystic area from dense tissue in choroid plexus organoids, and its application to other contexts or features, such as core darkening, remains to be tested.

Although selected to span highly distinct imaging scenarios, the testing dataset B used to benchmark different segmentation approaches comprised only ten brightfield images. As a consequence, the reported results provide only an approximate indication of the performance of each tool and larger annotated datasets would enable more comprehensive and standardized benchmarking.

Finally, the case studies were designed as proofs of concept and included a limited number of batches and conditions, limiting the biological interpretation of the findings reported. Additionally, except for the first case study, pixel-level ground-truth masks were not available. In those cases, segmentation quality was instead assessed by visual inspection of masks against raw images and through the quality-control outputs of Organotchi (number of organoids detected and number of outliers). While reducing the risk of segmentation errors propagating into the morphological analysis, this approach does not provide a quantitative measure of segmentation accuracy for these datasets.

## Resource availability

Organotchi will be made available on GitHub upon publication, together with installation and usage instructions and guidance on which segmentation route to select for different imaging and experimental settings, based on the benchmarking results reported here. The datasets used for case studies 2 and 4 (comprising the raw images and the corresponding masks generated with Organotchi) along with their respective metadata and the configuration files required to reproduce each analysis, will be deposited on Zenodo upon publication. All original analysis code will be made publicly available on GitHub upon publication.

## Acknowledgments

The authors thank Dr. Luca Deininger, Dr. Federico Colombo and Prof. Dr. Rasmus Schröder for their input and guidance throughout the course of this project; and Dr. Annamarija Raic and Yaping Li for providing the MCF-7 breast cancer spheroid dataset (case study 3) and their critical review of the manuscript. The authors used Claude (Opus 5; Anthropic) to improve the language and readability of the text. All content was subsequently reviewed and edited by the authors, who take full responsibility for the content of the publication. Icons in Figure 1 were created in BioRender (https://BioRender.com/38rlz1s).

## Author contributions

L.B. conceptualization, data curation, formal analysis, methodology, software, validation, writing - original draft preparation; T.L. software, writing - review & editing; D.A. conceptualization, experimentation, writing - review & editing; Z.Y. conceptualization, experimentation, writing - review & editing; F.C. experimentation, writing - review & editing; T.K. experimentation, writing - review & editing; I.P. experimentation, writing - review & editing; P.C. experimentation, writing - review & editing; K.P.S. experimentation, writing - review & editing; J.M. funding acquisition, writing - review & editing; R.M. funding acquisition, resources, supervision, writing - review & editing; S.M. conceptualization, funding acquisition, resources, supervision, writing - review & editing.

## Supplementary figures

**Supplementary Figure S1.**
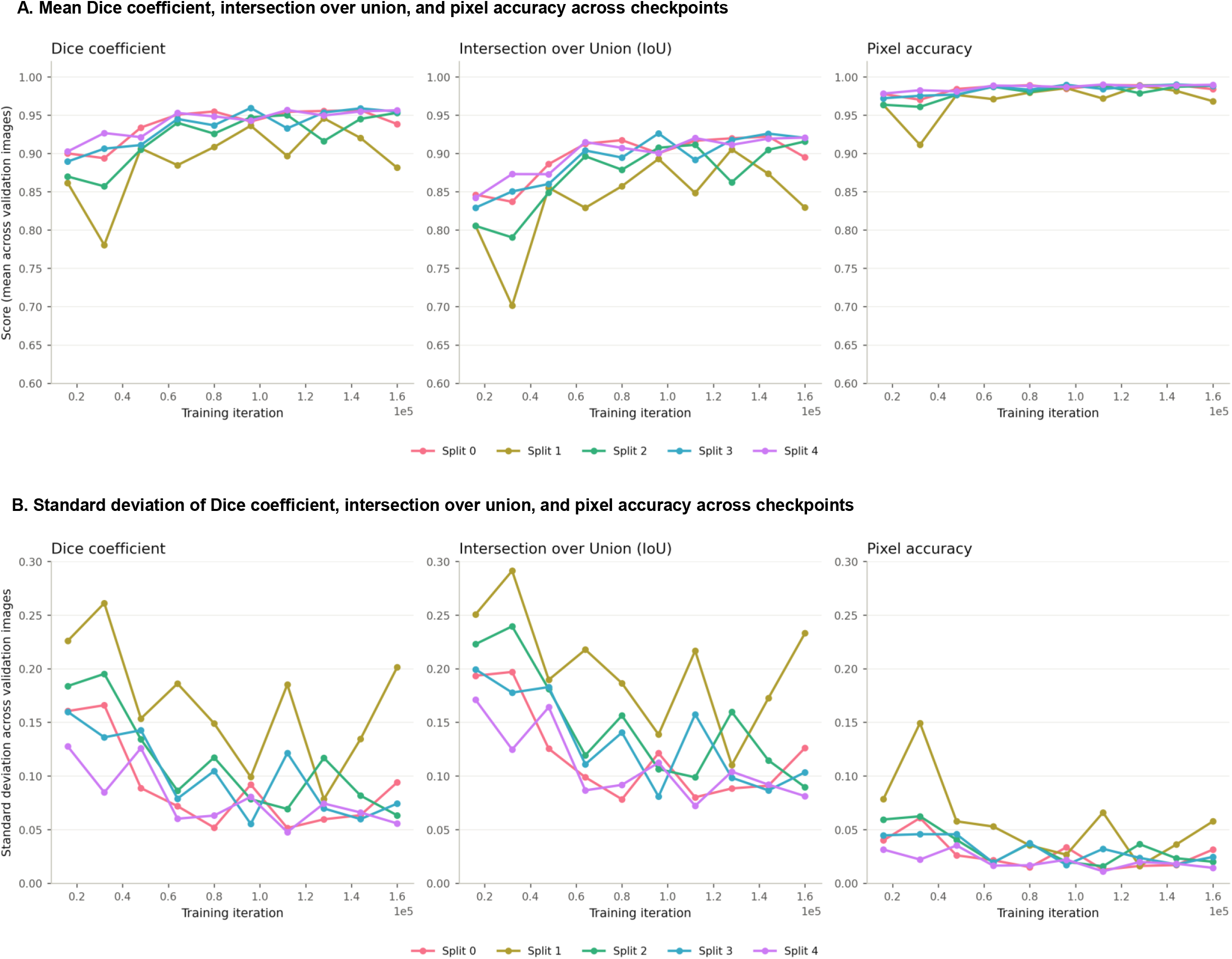
*SegFormer* performance across checkpoints. (A) Dice coefficient, Intersection over Union (IoU), and pixel accuracy, each as the mean across that checkpoint’s held-out validation images, plotted against training iteration (16,000-160,000, evaluated every 16,000 iterations) for each of 5 independent cross-validation splits (Splits 0-4). (B) The same three metrics’ standard deviation across held-out validation images, at the same checkpoints.

**Supplementary Figure S2.**
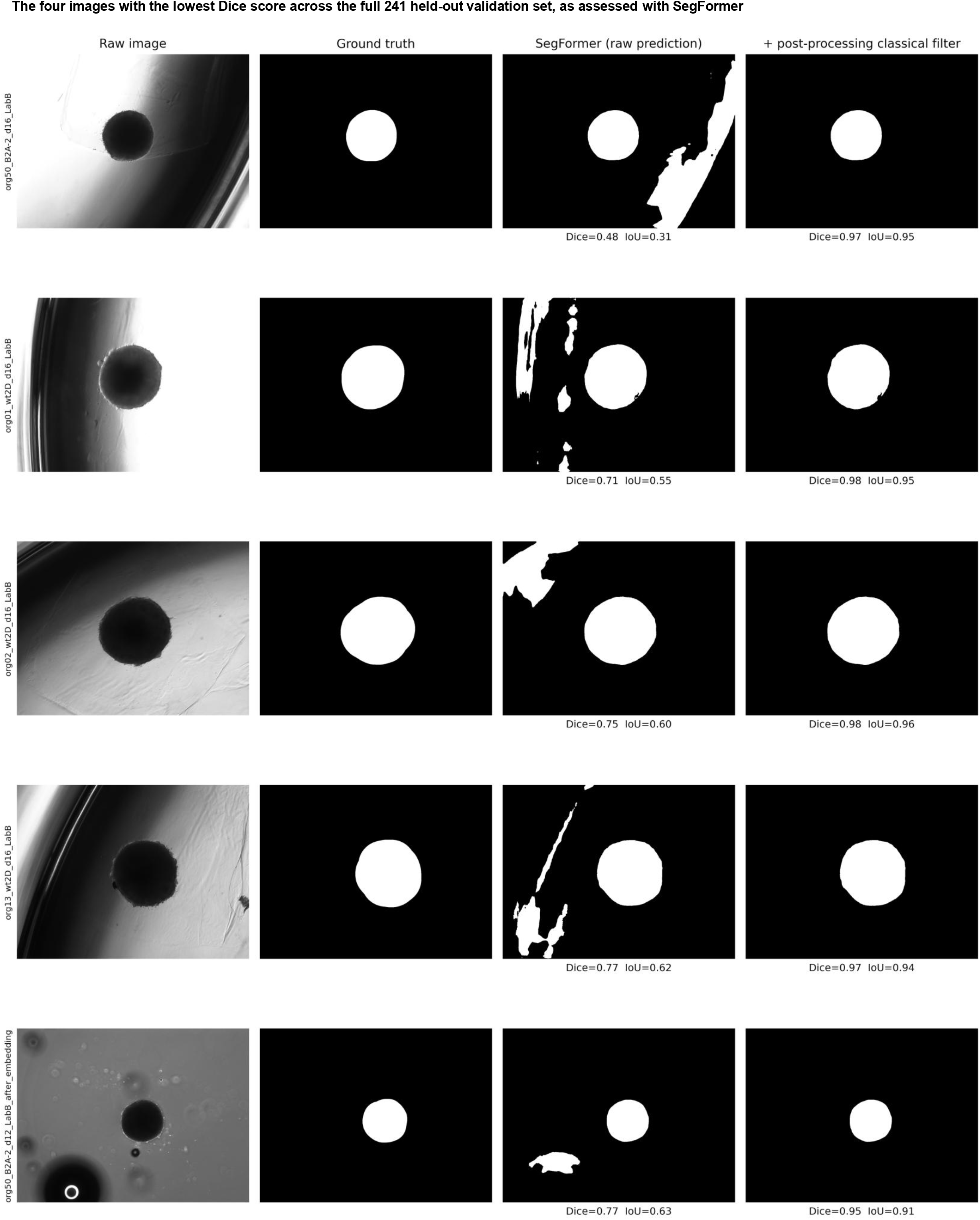
The five images with the lowest Dice score across the full 241 held-out validation set, as assessed with *SegFormer*. Each row of the figure presents, from left to right: raw image; ground truth mask; *SegFormer* prediction without and with an additional post-processing classical filter (see Methods).

**Supplementary Figure S3.**
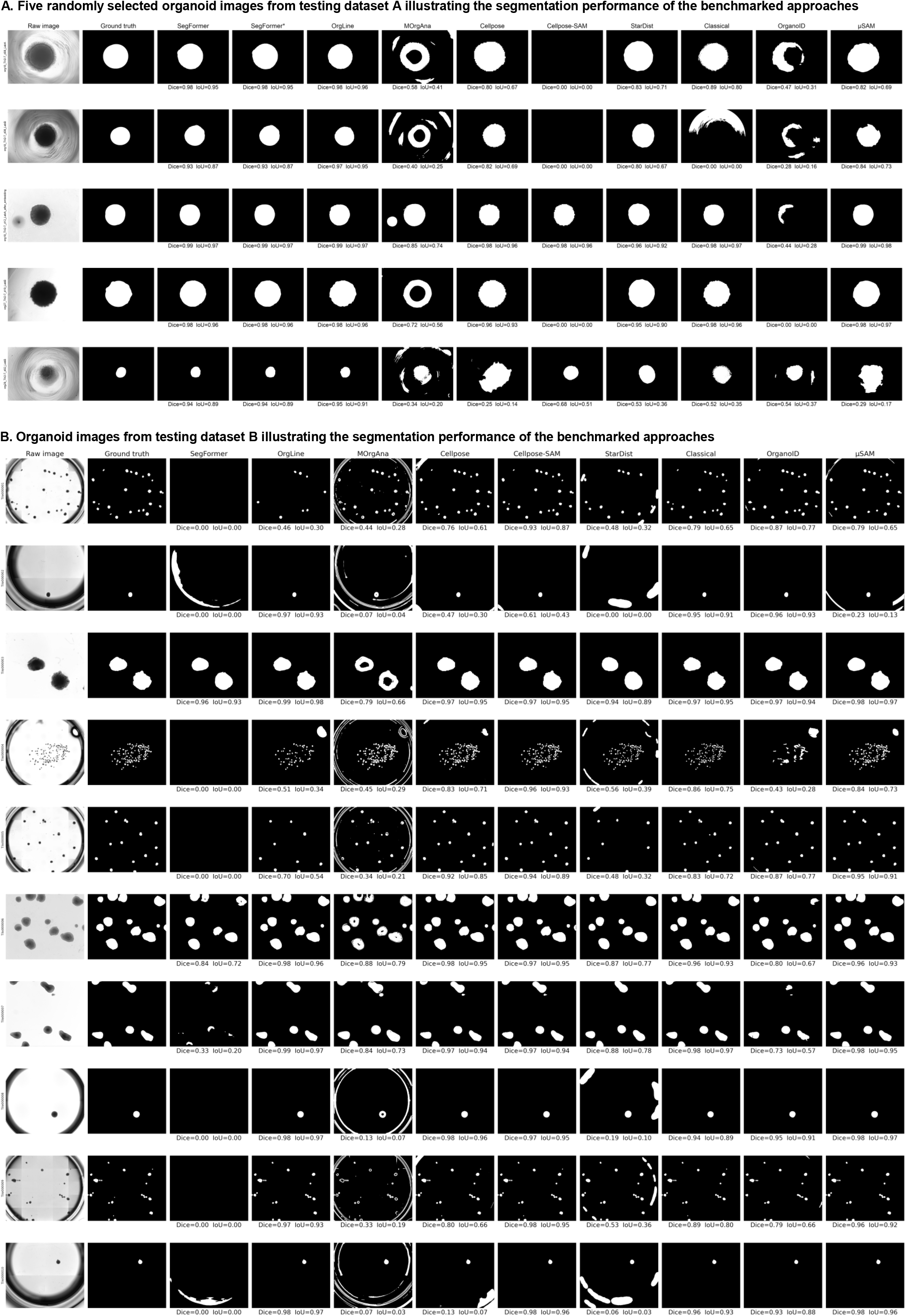
(A) Five randomly selected organoid images from testing dataset A illustrating the segmentation performance of the benchmarked approaches. Each row presents, from left to right: the raw image with its original label; the ground-truth mask; and the masks produced by the *SegFormer*-based model, the *SegFormer*-based model with an additional layer of classical post-processing filtering (*SegFormer*\*), *OrgLine*, *MOrgAna*, *Cellpose*, *Cellpose-SAM*, *StarDist*, *Classical*, *OrganoID*, and *μSAM*. Dice and Intersection over Union (IoU) values are shown beneath each mask. A summary of segmentation performance across all tools is presented in Figure 2A. (B) Similar to panel (A), but depicting the ten brightfield microscopy images from the testing dataset B and respective segmentation performance of the benchmarked approaches. A summary of segmentation performance across all tools is presented in Figure 2B.

**Supplementary Figure S4.**
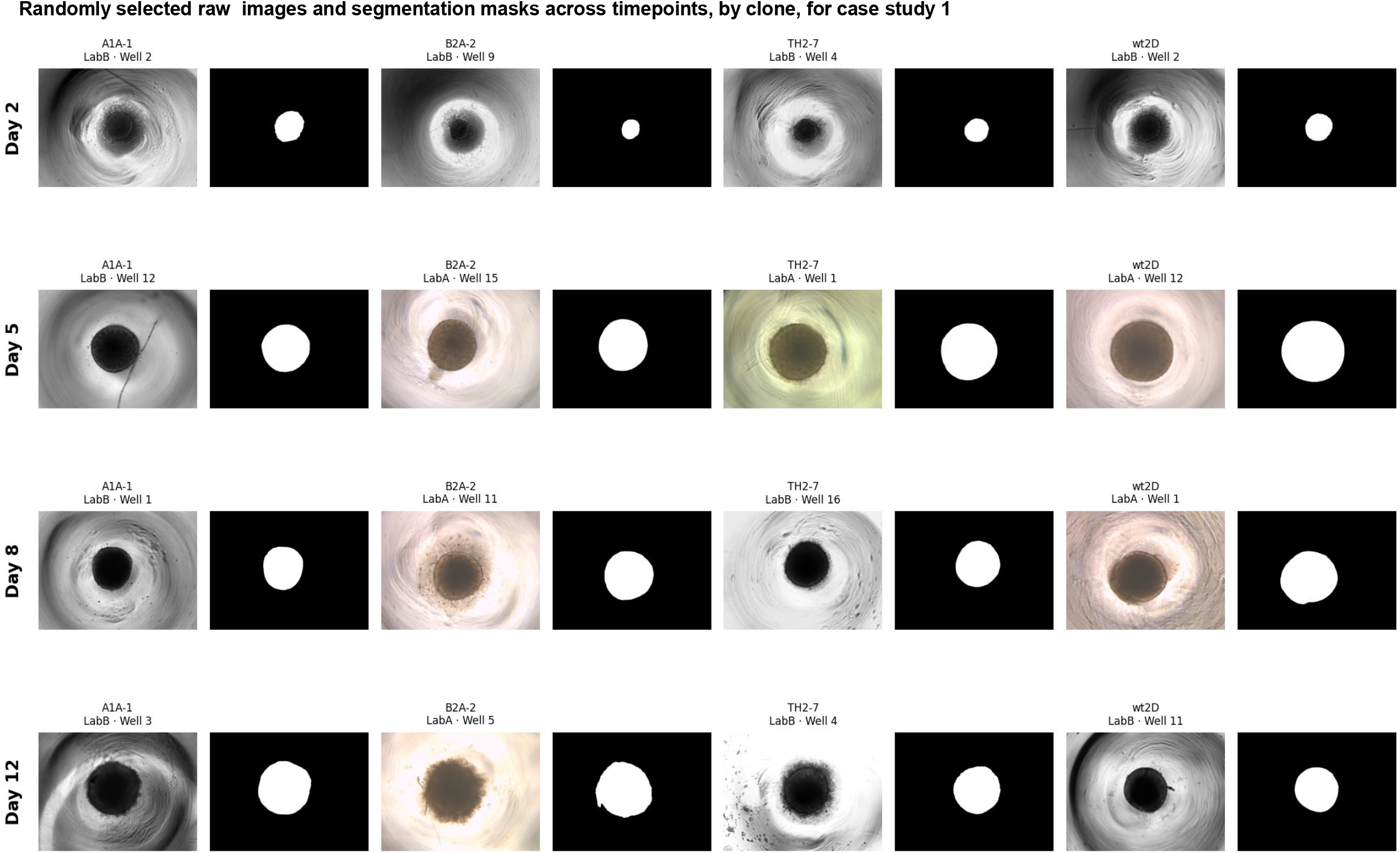
Randomly selected raw images and segmentation masks across timepoints, by clone, for case study 1: brain organoid morphology across four clones (A1A-1 = TUBA1A tubulinopathy; B2A-2 = TUBB2A tubulinopathy; TH2-7 = tyrosine-hydroxylase deficiency; wt2D = healthy control). Each column pair shows one experimental group’s raw image (left) and its corresponding predicted segmentation mask (right), labeled with the group, replicate/laboratory identity, and well number. Rows correspond to four imaging timepoints (Day 2, 5, 8, 12), earliest at top.

**Supplementary Figure S5.**
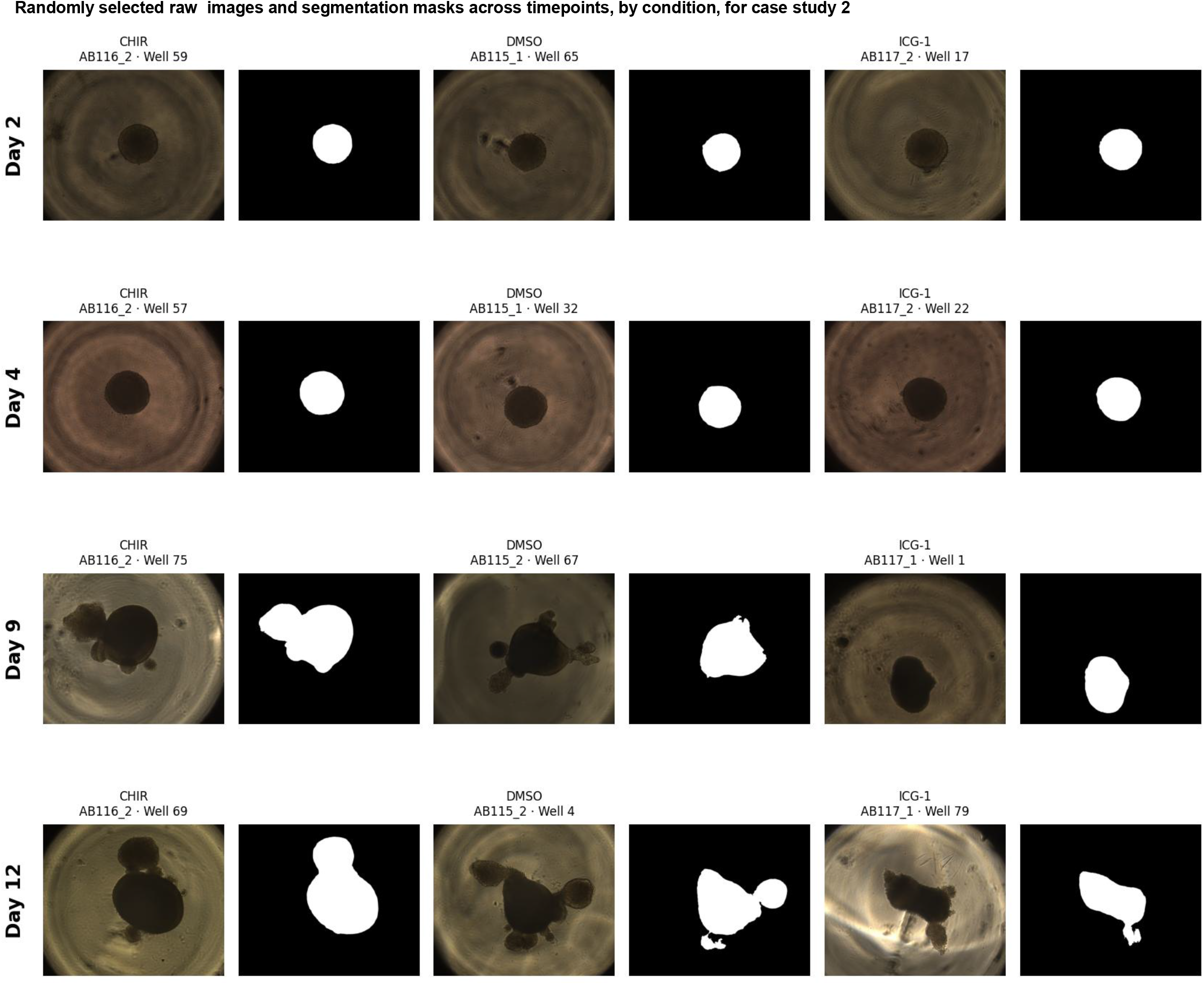
Randomly selected raw images and segmentation masks across timepoints, by condition, for case study 2: pharmacological modulation of Wnt signaling in dorsal forebrain organoids (CHIR: CHIR99021 = GSK3 inhibitor/Wnt activator; DMSO = vehicle control; ICG1: ICG-001 = beta-catenin/CBP inhibitor). Each column pair shows one experimental group’s raw image (left) and its corresponding predicted segmentation mask (right), labeled with the group, replicate/laboratory identity, and well number. Rows correspond to four imaging timepoints (Day 2, 4, 9, 12), earliest at top, chosen at random with a fixed seed.

**Supplementary Figure S6.**
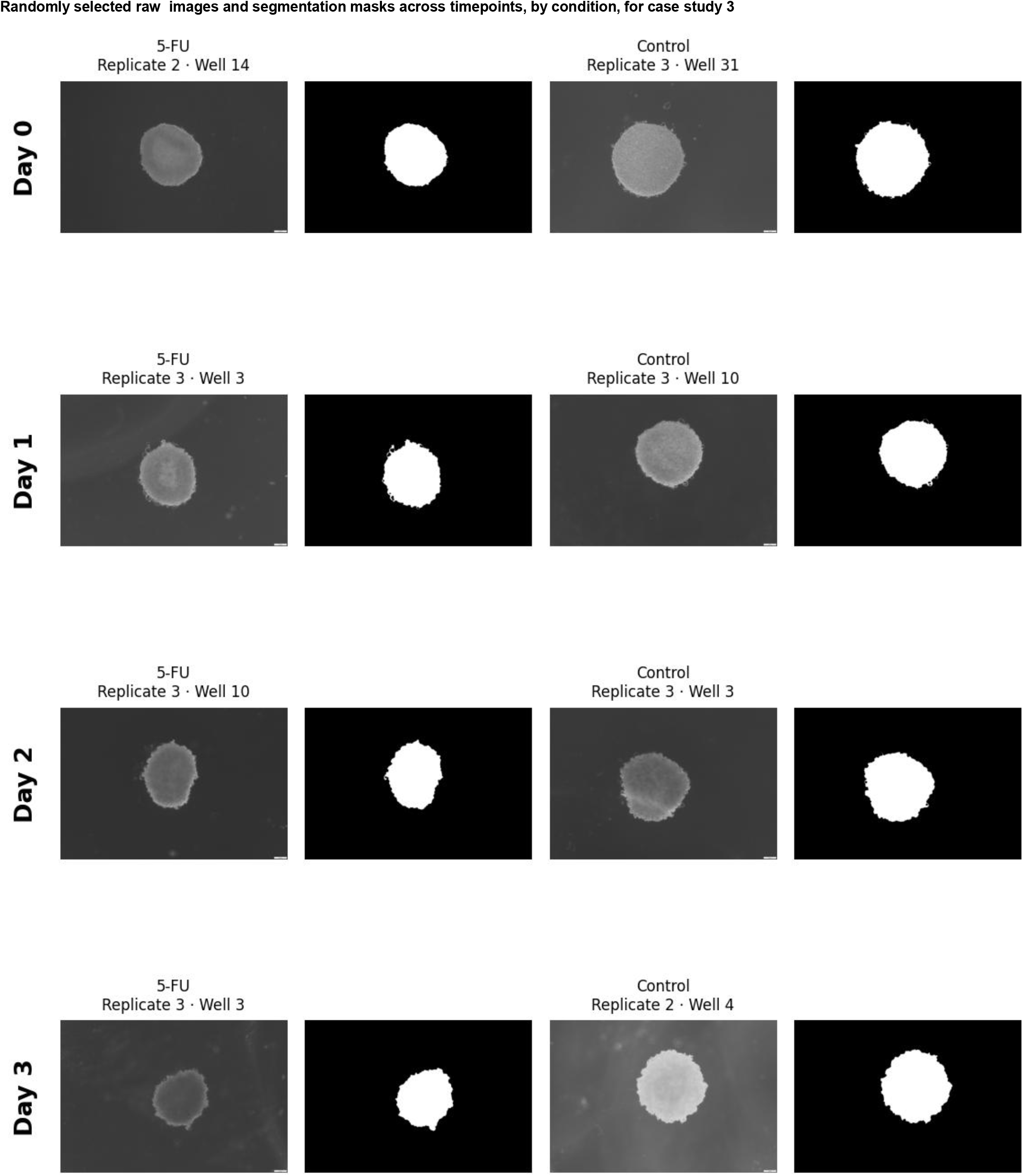
Randomly selected raw images and segmentation masks across timepoints, by condition, for case study 3: exposure response of MCF-7 spheroids to 5-FU vs. Control, tracked across days 0-3. Each column pair shows one experimental group’s raw image (left) and its corresponding predicted segmentation mask (right), labeled with the group, replicate/laboratory identity, and well number. Rows correspond to four imaging timepoints (Day 0, 1, 2, 3), earliest at top, chosen at random with a fixed seed.

**Supplementary Figure S7.**
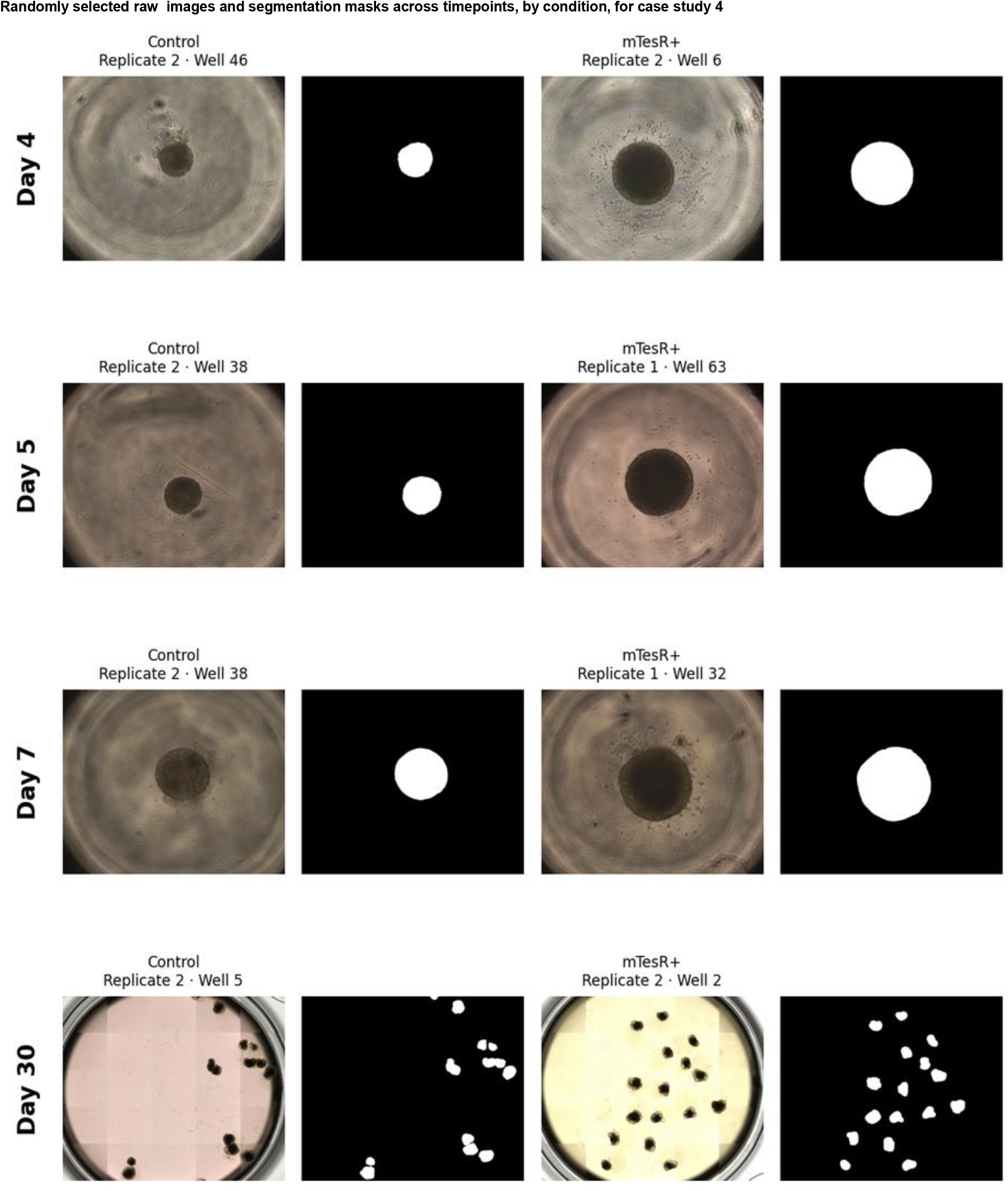
Randomly selected raw images and segmentation masks across timepoints, by condition, for case study 4: Effects of media composition in choroid-plexus organoid differentiation (mTesR Plus vs. Control). Each column pair shows one experimental group’s raw image (left) and its corresponding predicted segmentation mask (right), labeled with the group, replicate/laboratory identity, and well number. Rows correspond to four imaging timepoints (Day 4, 5, 7, 30), earliest at top, chosen at random with a fixed seed.

**Supplementary Figure S8.**
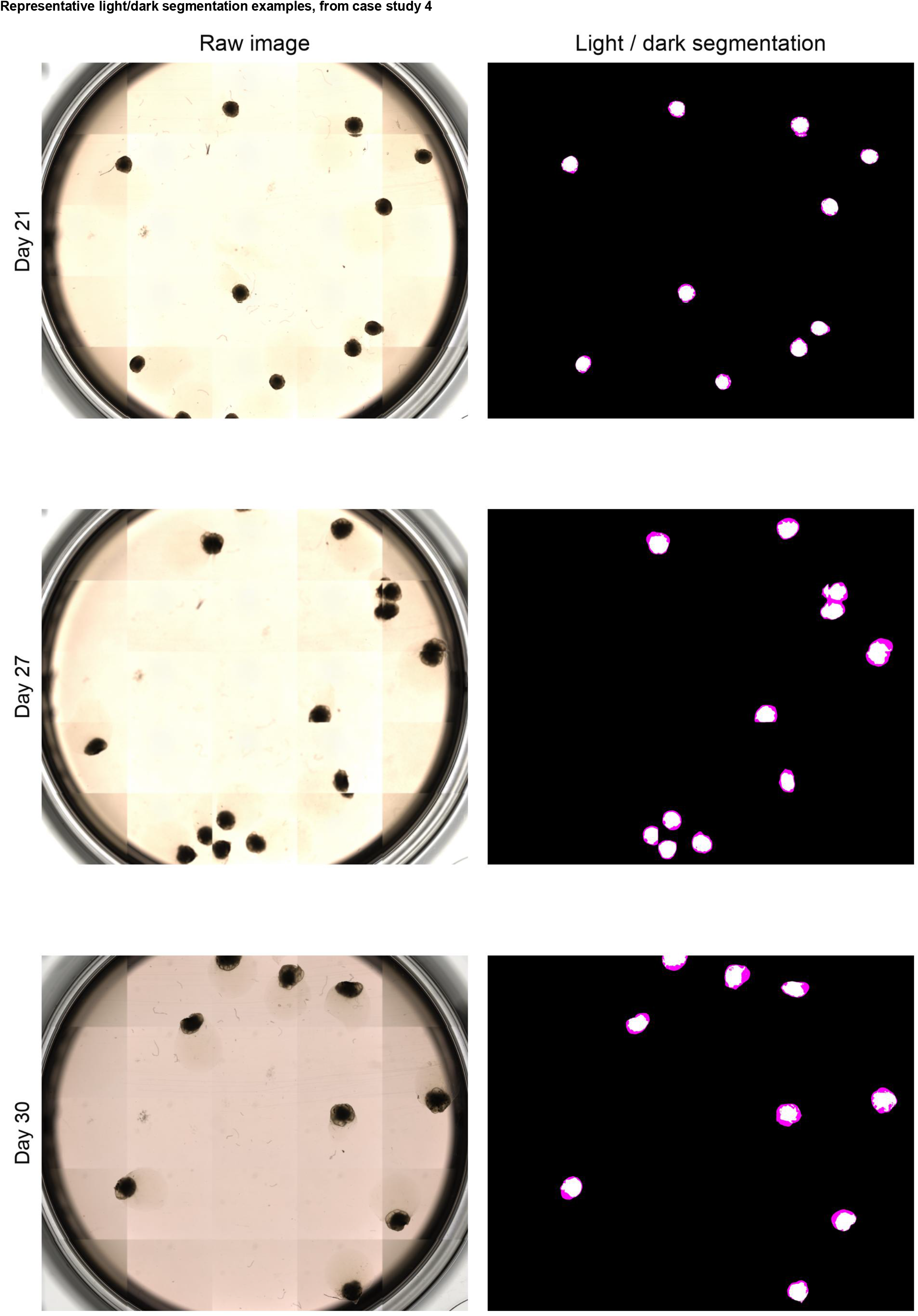
Representative light/dark segmentation examples, from case study 4. Raw brightfield image (left) and the corresponding per-organoid light/dark overlay (right; black = background, white = each organoid’s own darker “dark” region, magenta = its lighter “light” region, from an Otsu brightness split of each organoid’s own mask; see Methods), for one representative well at days 21, 27, and 30.

**Supplementary Figure S9.**
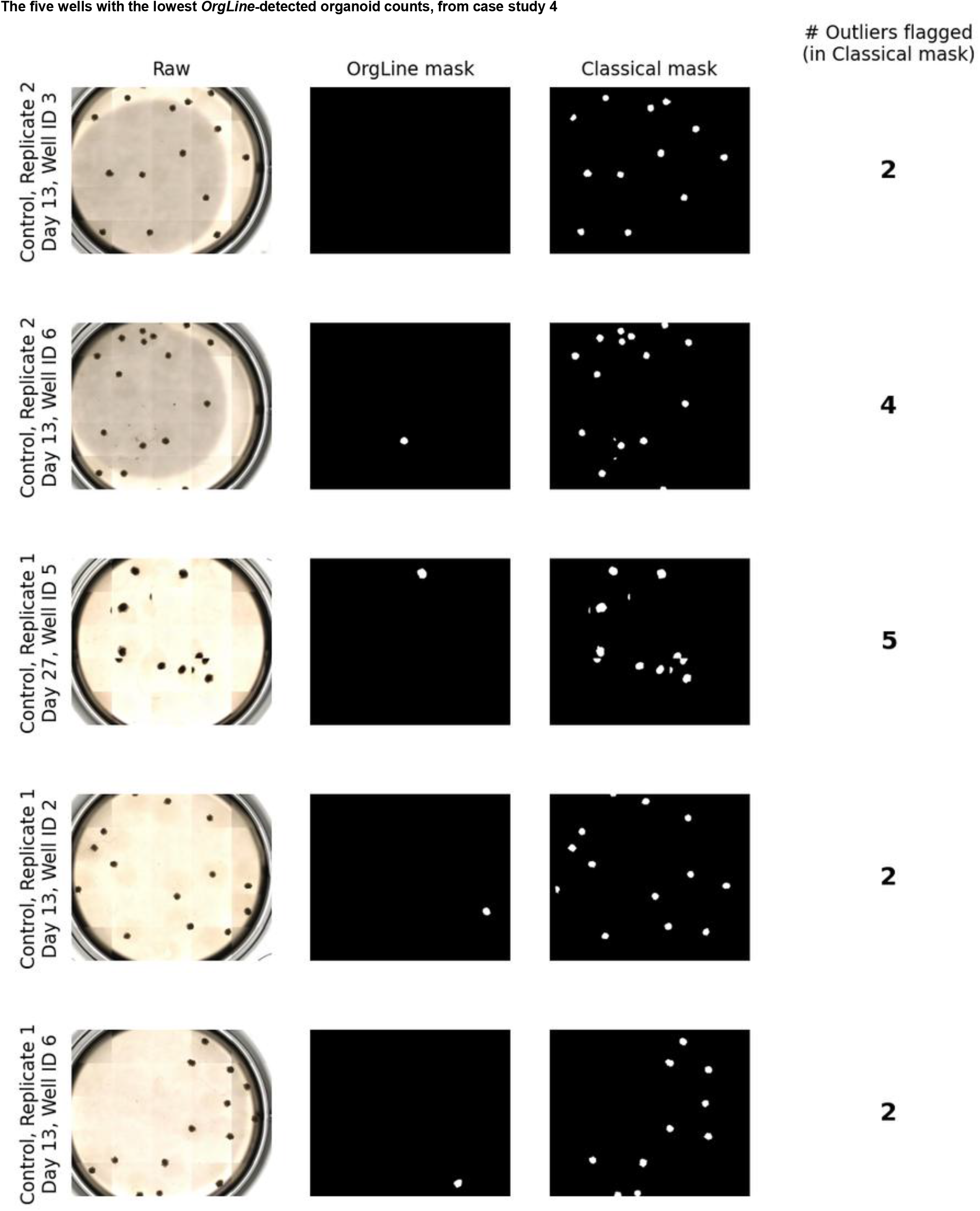
The five wells with the lowest *OrgLine*-detected organoid counts, from case study 4 (from day 13 onwards; ties in the number of detected objects broken at random with a fixed seed). Each row shows, left to right: the raw (stitched) well image; *OrgLine*’s predicted mask; the obtained mask using the *Classical* segmentation approach; and the number of objects flagged as outliers (Tukey IQR fencing) while using the *Classical* segmentation approach.

**Table S1.** Performance of every segmentation tool/approach evaluated in this manuscript, restricted to 264 brightfield microscopy images from the Schröter *et al*, 2024 dataset (testing dataset A) and an *ad hoc* dataset of 10 brain organoid images spanning a range of magnifications, single- and multi-organoid fields of view, and several regional identities and morphologies (testing dataset B). Each row reports one segmentation approach on one dataset (A or B), giving the mean, standard deviation, median, and 25^th^/75^th^ percentiles for Dice coefficient, Intersection over Union (IoU), pixel accuracy, predicted and ground-truth object counts, Average Precision at IoU 0.50 and 0.75 (AP50, AP75), mean AP across IoU thresholds 0.50-0.95 (mean AP), and F1 score at IoU 0.50 (F1@50). n_images denotes the number of images each row’s statistics are computed over.

For each case study, we provide 3 tables (A-C) that complement the figures for each of the case studies on the morphological analyses performed. **Tables S2A-S2C** correspond to case study 1, **Tables S3C-S3C** to case study 2, **Tables S4A-S4C** to case study 3, and Tables **S5A-S5C** to case study 4. Since the structure of each of the 3 tables (A-C) is identical across case studies, we provide a single legend for each below, from A to C.

**A.** Day-by-day pairwise statistical comparisons for every morphological feature and every grouping variable defined for this case study, at each of its designated comparison days. One row per (feature, grouping variable, day, pair of groups): Welch’s t-test on well-level means, with Hedges’ g effect size; p-values are Benjamini-Hochberg FDR-corrected across all comparisons in this table.

**B.** Morphological features distinguishing each HDBSCAN cluster from every other clustered organoid on the same day. One row per (cluster, feature): cluster and rest-of-dataset means, Welch’s t-test/Hedges’ g, and Benjamini-Hochberg FDR-corrected significance across every cluster-by-feature test for that clustering run.

**C.** Outlier rate (Tukey interquartile-range rule, 1.5× fence) for each morphological feature, condensed to one row per feature: the mean outlier percentage across every group-by-day cell, and the single group/day combination with the highest outlier percentage for that feature, together with its value.

## Methods

### Training, validation and testing of *SegFormer*-based semantic segmentation model

The *SegFormer*-based segmentation model incorporated in Organotchi was trained on the published dataset of Schröter *et al*, 2024, which includes approximately 1,400 brightfield images and matched ground-truth masks from 64 tracked organoids, generated from 4 clones and 2 imaging laboratories. Model training, validation, and testing were conducted using the MMSegmentation 1.2.2 framework (PyTorch 2.1.0, MMEngine 0.10.7, MMCV 2.1.0, MMDetection 3.3.0), employing *SegFormer* with the smallest implemented encoder (MiT-B0) as the semantic segmentation backbone. The model was trained using the AdamW optimizer (learning rate = 0.00006, β₁ = 0.9, β₂ = 0.999, weight decay = 0.01) with a batch size of 1 for 160,000 iterations. Validation was performed every 16,000 iterations. The loss function was a weighted combination of binary cross-entropy and Dice loss (1:10 ratio). On-the-fly data augmentation included (1) random resizing, (2) random cropping, (3) random horizontal flipping with a 0.5 probability, and (4) image distortion. The model was trained independently on 5 cross-validation splits, each using independent ≈63%/17%/20% train/validation/test partitions of the dataset.

Segmentation performance was evaluated on each checkpoint and for each cross-validation split through the calculation of pixel-level Dice coefficient, intersection-over-union (IoU), and pixel-wise accuracy on the binarized organoid-versus-background mask. For a predicted binary mask P and its corresponding ground-truth binary mask G (organoid pixels = 1, background = 0, both defined over the same image of N total pixels), Dice was computed as

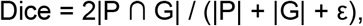

IoU as

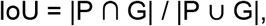

and pixel-wise accuracy as the fraction of pixels at which P and G agree,

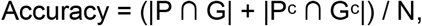

where |·| denotes pixel count, Pᶜ/Gᶜ denote the background (complement) sets, and ε = 10⁻⁶ is a small constant added to Dice’s denominator for numerical stability.

The model achieving the highest mean Dice coefficient minus one standard deviation of the five splits on the respective validation subset (mDice - *k* × Dice Std, *k* = 1; split 4’s iteration-112,000, with mDice 0.957 and Dice Std 0.0477) was selected and adopted throughout this study for every *SegFormer*-based segmentation run described in this manuscript.

### Classical segmentation approach

Organotchi offers a fully classical segmentation approach based on the scikit-image 0.25.2 and opencv-python 4.10.0.84 packages, consisting of the following processing steps: (1) the raw image is first converted to a binary foreground/background mask using a single global grayscale threshold (dark-foreground polarity, with a default cut-off at 128 on an 8-bit image), or the Otsu thresholding algorithm; (2) an optional central circular mask (disabled by default) restricts the analysis to a given fraction of the image width; (3) an optional morphological closing operation (disabled by default) bridges small gaps in an incompletely thresholded object boundary; (4) foreground pixels are labeled into connected-component candidate regions (using 8-connectivity) and filtered by minimum and maximum area, and minimum circularity (computed using 4 × π × Area / Perimeter²);(5) for images expected to contain a single organoid, only the largest surviving contour by area is retained; for multi-organoid images, all surviving contours are retained. All steps/filtering parameters and respective thresholds used are user-customizable. Default minimum circularity is set at 0.2 and default area values should fall in 0.03-3.0 mm² range.

### OrgLine-based segmentation approach

As an additional segmentation backend alongside Organotchi’s previously described SegFormer-based and classical approaches, we integrated *OrgLine* 1.0.0 (Deng *et al*, 2026), using its smallest published (“tiny”) detector/encoder checkpoint. When employed, OrgLine’s detections were kept above a confidence threshold of 0.2 and de-duplicated by non-maximum suppression at an IoU threshold of 0.7; both values are user-customizable. For images expected to contain a single organoid, only the highest-confidence detection is retained; for multi-organoid images, all detections surviving the confidence and IoU thresholds are kept.

### Optional post-processing filter

An optional post-processing filter can be applied on top of the *SegFormer*- and the *OrgLine*-based segmentation approaches to remove potential imaging artifacts. This post-processing filter consists of the following steps: (1) removal of objects below a minimum area (using skimage.measure.regionprops, 8-connectivity; default set at 0.01 mm²); (2) optional erosion (disabled by default); (3) removal of objects below a minimum circularity (computed using 4 x π x Area / Perimeter²; default 0.1); (4) if more objects than expected survive, only the most circular one(s) are kept; optional dilation and hole-filling (default disabled).

This post-processing filter was applied in all brain organoid case studies for the period during which organoids were imaged at 4-5× magnification in 96-well plates.

### Benchmarking against existing segmentation tools

We evaluated segmentation accuracy of Organotchi’s three available segmentation approaches, together with a set of published organoid- and cell-scale instance segmentation tools, using two distinct brain datasets. The first dataset (testing dataset A) comprised the 264 brightfield, single-organoid-per-well images assigned to the held-out test partition of the selected SegFormer-based checkpoint’s cross-validation split (split 4), together with their published ground-truth masks (Schröter et al., 2024). The second dataset (testing dataset B) consisted of 10 iPSC-derived organoid images imaged with ECHO Spinning Disk Confocal microscope at 1.25× magnification. The respective ground truth masks were produced manually in napari 0.8.0, with every visually distinct organoid delineated as a separate instance. Additional information concerning the differentiation protocols, imaging settings, and culture strategies used for each image is listed on the table below.

| **Tile ID** | **Regional identity** | **Organoid differentiation protocol based on** | **Differentiation day** | **Number of organoids per plate** | **Tile grid** | **Image dimensions** |
|---|---|---|---|---|---|---|
| Tile000001 | Choroid Plexus | Pellegrini *et al*, 2020 | 13 | Multiple | 5×5 | 11952 px × 9956 px |
| Tile000002 | Cerebellum | Atamian *et al*, 2024 | 32 | Single | 2×2 | 4800 px × 4001 px |
| Tile000003 | Choroid Plexus | Pellegrini *et al*, 2020 | 24 | Multiple | 1×1 | 2416 px × 2016 px |
| Tile000004 | Cerebellum | Atamian *et al*, 2024 | 21 | Multiple | 5×5 | 11952 px × 9956 px |
| Tile000005 | Choroid Plexus | Pellegrini *et al*, 2020 | 13 | Multiple | 5×5 | 11952 px × 9956 px |
| Tile000006 | Cerebellum | Atamian *et al*, 2024 | 22 | Multiple | 1×1 | 2416 px × 2016 px |
| Tile000007 | Cerebellum | Atamian *et al*, 2024 | 19 | Multiple | 1×1 | 2416 px × 2016 px |
| Tile000008 | Cerebellum | Atamian *et al*, 2024 | 70 | Single | 2×2 | 4800 px × 4001 px |
| Tile000009 | Cerebellum | Atamian *et al*, 2024 | 33 | Multiple | 5×5 | 11952 px × 9956 px |
| Tile000010 | Cerebellum | Atamian *et al*, 2024 | 32 | Single | 2×2 | 4800 px × 4001 px |

Apart from this pipeline’s classical segmentation, *SegFormer*-based (both without and with the post-processing filter), *OrgLine*-based segmentation approaches (all as described above), we compared against: *Cellpose* (cyto3 pretrained checkpoint, cellpose 3.1.1.3); *Cellpose-SAM* (cpsam pretrained checkpoint, cellpose 4.2.1.1); *StarDist* 0.9.2 (2D_versatile_he pretrained model, on RGB input, at default detection thresholds); *MOrgAna* 0.1.2 (Python 3.9; trained on 5 images sampled from this split’s own held-out validation partition — the same partition used for *SegFormer* checkpoint selection, and by construction disjoint from the test partition reported here); *OrganoID* (bundled pretrained TensorFlow-Lite model, default detection threshold); and *μSAM* 1.8.8 (vit_b_lm pretrained checkpoint, run in Automatic Instance Segmentation mode).

As segmentation accuracy of *StarDist*, *Cellpose*, *Cellpose-SAM*, and *μSAM*, is sensitive to an expected object-scale parameter (diameter for *StarDist*, *Cellpose*, and *Cellpose-SAM* and minimum instance area for *μSAM*), we supplied per-image estimates for each tool, since organoid size changes substantially from image to image (depending, for example, on the culture period, and magnification used). Per-image estimates were derived from *OrgLine*’s own predicted instance sizes for the same image, given its overall high segmentation accuracy in the considered datasets.

Every tool was run on CPU-only hardware in its own isolated software environment, following that tool’s own installation documentation; the *SegFormer*-based model, *OrgLine*, and the general Organotchi pipeline were developed and implemented separately (PyTorch 2.1.0, Python 3.10).

For each tool, segmentation accuracy was evaluated through the calculation of two families of metric. Pixel-level Dice coefficient, intersection-over-union (IoU), and pixel-wise accuracy were computed on the binarized organoid-versus-background mask, independent of any instance separation. Instance-level accuracy was evaluated via average precision (AP) at ten IoU thresholds from 0.50 to 0.95, in steps of 0.05. At each threshold t, predicted and ground-truth instances were matched one-to-one by a greedy, highest-IoU-first procedure, run separately at every threshold. Matched pairs were counted as true positives (TP); predicted instances left unmatched, as false positives (FP); ground-truth instances left unmatched, as false negatives (FN). AP at threshold t was then defined as

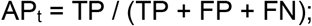

F1 at threshold t was computed from the same TP/FP/FN as

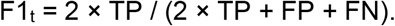

AP at t=0.50 (AP50) and F1 at t=0.50 (F1@50) were reported for each tool.

Predicted instances of semantic models (*SegFormer* and *MOrgAna*) were derived from their binary output by connected-component labelling. Two touching organoids correctly segmented at the pixel level but merged into one connected region are therefore counted as a single predicted instance against two ground-truth instances for these methods.

### Cell culture, spheroid formation and organoid differentiation

For case study 1, iPSC culture and organoid differentiation is described in the original publication (Schröter *et al*, 2024). Similarly, for case study 3 cell culture and spheroid formation is described in Raic *et al*, 2026. Here we describe iPSC culture and organoid differentiation for case studies 2 and 4.

#### Case study 2: Pharmacological modulation of Wnt signaling in dorsal forebrain organoids

The commercially available human iPSC line BIONi010-C (Source: EBiSC; RRID: CVCL_1E68) was used for this experiment. The iPSCs were maintained in mTeSR Plus medium (STEMCELL Technologies, Cat. no. 100-0276) at 37 °C in a humidified atmosphere containing 5% CO_2_ Cells were passaged onto hESC-qualified growth factor-reduced Matrigel-coated (Corning, Cat. no. 354277) using Gentle Cell Dissociation Reagent (STEMCELL Technologies, Cat. no. 07174). After passaging media was supplemented with 2 μM thiazovivin (Sigma, Cat. no. 420220) and replaced with thiazovivin-free fresh media the next day. All cell lines were tested for pluripotency with antibodies against OCT4 (rabbit, 1:500, Abcam, Cat. no. ab19857) and mycoplasma contamination (Abcam, Cat. no. ab289834) upon thawing. Cells with passage number below 20 were used for experiments.

Dorsally patterned forebrain organoids were generated by using previously described protocol (Velasco *et al*, 2019) with minor modifications. iPSC cultures at confluency level about 80-90% were treated with 10 µM Y-27632 (Cayman Chemical, Cat. no 10005583) for 1 hour prior to dissociation. Then, cells were dissociated using Accutase (Merck, Cat. no. A6964) to obtain single-cell suspension, and 9,000 cells were seeded per well in 96-well ultra-low attachment plates (S-bio, Cat. no. MS-9096VZ) in mTeSR Plus media supplemented with 10 µM Y-27632.

The following day, designated as day 1, aggregates were checked and the medium was replaced with cortical differentiation media (CDM), consisted of Glasgow’s Minimum Essential Medium (GMEM, Gibco, Cat. no. 11710035) with 20% KnockOut Serum Replacement (KOSR, Thermofisher Scientific, Cat. no. 10828028), 1X MEM non-essential amino acids (Sigma, Cat. no. M7145), 0.11 mg/mL Sodium Pyruvate (Thermofisher Scientific, Cat. no. 11360070), 1X Penicillin-Streptomycin (Sigma, Cat. no. P0781), 0.1 mM 2-Mercaptoethanol (Gibco, Cat. no. 21985023) with supplements 20 µM Y-27632, 3 µM IWR (Merck, Cat. no. 681669), and 5 µM SB431542 (Merck, Cat. no. 616461).

Organoids were assigned to one of the three treatments conditions: vehicle - 1:1000 DMSO (Fisher, Cat. no. 10103483), CHIR - 3 µM CHIR-99021 (Tocris, Cat. no. 4423), or ICG-001 - 3 µM ICG-001 (Tocris, Cat. no. 4505). On day 4, the medium was replaced with freshly prepared CDM containing the same supplements. On days 7 and 10, the medium was replaced with freshly prepared CDM supplemented with IWR-1, SB431542, and the corresponding treatment compound.

#### Case study 4: Effects of media composition in choroid-plexus organoid differentiation

The commercially available human iPSC line BIONi010-C (Source: EBiSC; RRID: CVCL_1E68) was used for this experiment. The iPSC line was cultured under standard conditions (37°C, 5% CO_2_, and 100% humidity) and the respective cells were maintained in mTESR Plus media and passaged onto hESC-qualified growth factor-reduced Matrigel-coated plates in colonies using Gentle Dissociation Reagent.

STEMdiff™ Choroid Plexus Organoid Differentiation Kit and STEMdiff™ Choroid Plexus Organoid Maturation Kit were used for choroid plexus organoid generation and culturing. Stem Cell Technologies choroid plexus differentiation protocol was followed with minor modifications (STEMCELL^TM^ Technologies; Pellegrini *et al*, 2020). Instead of U-shaped 96 well plates, ultra-low attachment V-shaped 96 well plates were used. For this case study, a batch of iPSCs was seeded in iPSC media (mTESR Plus) and another directly in the kit differentiation media (Control). Organoids were kept in 96-well plates until Matrigel embedding. After Matrigel embedding, they were put on the shaker inside the incubator on day 10. The shaker settings were: 37°C, 5.0% CO_2_ and 56 RPM (eppendorf New Brunswick S41i incubator). The remaining media changes with specific media requirements were performed as described in the original protocol (STEMCELL^TM^ Technologies)..

### Image acquisition for the reported case studies

For case studies 1 and 3, image acquisition is described in the original publications (Schröter *et al*, 2024; Raic *et al*, 2026). For the remaining case studies (2 and 4), organoids were imaged using an ECHO Spinning Disk Confocal microscope. When in 96-well plates, organoids were imaged with a 4× magnification and a 1×1 tile grid. When in 6-well plates, organoids were imaged with a 1.25× magnification and a 5×5 tile grid.

### Image pre-processing

Tile stitching is performed by the Organotchi’s pipeline itself from raw, unstitched tiles, producing sequentially-numbered stitched images (Tile000001.jpg, …) that all downstream segmentation steps treat as the unit of input. 16-bit source TIFFs are rescaled to 8-bit by linear min-max normalization. During stitching, this normalization is computed jointly across every tile belonging to the same stitched image.

### Morphology feature extraction and statistical comparisons across conditions

Physical pixel size was calibrated per microscopy magnification and image resolution used in each case study. For case studies 1 and 3, calibration values adopted from the respective original publications (Schröter *et al*, 2024; Raic *et al*, 2026) (case study 1: 0.316 × 0.316 μm/px for Lab A and 0.334 μm/px × 0.334 μm/px for Lab B; case study 3: 0.869565 μm/px × 0.869565 μm/px). For the remaining case studies (2 and 4), a pixel size of 0.888 × 0.887 μm/px was used for imaged acquired at 4× magnification, and 2.842 × 2.839 μm/px for images acquired at 1.25× magnification.

Well identity (plate and well index) was recovered from each stitched image’s sequential tile number and the number of images expected per plate for that organoid batch and run (imaging day). For every segmented organoid instance, ten morphological features were computed from its labeled mask region (skimage.measure.regionprops, 8-connectivity): area (μm²); diameter (μm, computed as sqrt(4×Area/π)); perimeter (μm); maximum Feret diameter (μm); aspect ratio (major axis length/minor axis length of the best-fit ellipse of equivalent second moments); circularity (4×π×Area/Perimeter²); roundness (4×Area/(π×maximum Feret diameter²)); solidity (area / convex-hull area); eccentricity (of the best-fit ellipse); and skeleton length (μm, the pixel count of the morphological skeleton of the object’s own cropped mask, converted to physical units).

Area was converted from pixels to μm² using the true, anisotropic (x, y) pixel spacing. All other length measurements (perimeter, Feret diameter, skeleton length) used the average of the x and y spacings.

Two levels of aggregation are used throughout for statistical comparison and general morphological characterization across groups: “object-level” data or “well-level” data (used wherever a comparison is explicitly described as being tested on well means, to avoid pseudoreplication from unequal numbers of organoids per well). These two aggregation levels correspond to the unweighted arithmetic mean of each feature across every organoid detected in that image/well on that day.

Outliers were flagged, and optionally removed prior to any other analysis, by Tukey fencing (values beyond 1.5× the interquartile range from the first/third quartile), computed on log-transformed values for the five strictly-positive, right-skewed size features (area, diameter, perimeter, maximum Feret diameter, skeleton length) and on raw values for the five dimensionless shape descriptors (aspect ratio, circularity, roundness, solidity, eccentricity). This exclusion step was applied to individual organoid measurements (“object-level”).

Coefficient of variation was computed as 100% × (standard deviation / mean) of a feature within a group and timepoint.

Between-group comparisons at a single timepoint used a two-sided Welch’s t-test (SciPy 1.15.3; unequal variances assumed) and Hedges’ g effect size, computed on well-level means unless otherwise noted. P-values were corrected for multiple comparisons using the Benjamini-Hochberg false-discovery-rate procedure (FDR).

Longitudinal trends were summarized as the group mean at each timepoint +/− the standard error of the mean. Optionally, the standard deviation or a parametric 95% confidence interval can also be provided.

Per-well growth rate, k, was estimated by fitting an exponential growth model (NumPy 1.26.4), Area(t) = Area_0_×exp(k×t), via ordinary linear regression of log(feature) on day, to each physical well’s own time course (at least 3 timepoints required); doubling time was reported as ln(2)/k. For wells imaged in a multiple-organoids-per-image regime, where individual organoids have no stable identity from one day to the next, the value fitted for a given day was the mean across every organoid detected in that well’s image that day, so that a well-level growth rate could still be estimated despite the lack of organoid-level tracking.

Unsupervised structure was examined by projecting each day’s pooled, z-scored morphology features onto two dimensions by UMAP (umap-learn 0.5.12; 15 nearest neighbours, minimum distance 0.1) and by PCA, each run independently per day at the object level, using a non-redundant feature subset selected once per case study by pooling every analysed day’s data together. Starting from a fixed, ordered list of the 10 computed morphological features, each feature was retained unless its absolute Spearman correlation with an already-retained feature exceeded 0.9, in which case it was dropped in favour of that earlier, retained feature. Clusters were identified directly on each day’s own two-dimensional UMAP embedding using HDBSCAN (scikit-learn 1.7.2; minimum cluster size 5, excess-of-mass cluster selection). Points not confidently assigned to any cluster were excluded from downstream cluster-level statistics. Cluster identity was compared against each project’s experimentally-defined grouping variable(s) using the Adjusted Rand Index and Normalized Mutual Information (chance-corrected and uncorrected agreement, respectively, computed excluding unassigned points), weighted cluster purity, and a chi-square test of independence between cluster assignment and group label. The features distinguishing each cluster from every other clustered organoid that day were identified by Welch’s t-test/Hedges’ g per feature, Benjamini-Hochberg corrected across every cluster-by-feature test for that clustering run.

Light:dark sub-region within the organoids was computed from their own masked pixel values using an Otsu threshold (skimage.filters.threshold_otsu). Pixels darker than this intra-object threshold were classed as dark, the remainder as light. Separability of the resulting two-class split was quantified as eta-squared and a split was flagged as bimodal only once eta-squared reached a minimum value (default 0.3). Light and dark pixel counts were converted to area (μm²) via the respective pixel-to-micron calibration and used to report, per organoid: light area, dark area, light fraction (light area / total area), dark fraction (1 - light fraction), and light:dark ratio (light area / dark area, undefined when dark area is zero), together with the count and mean, median, and maximum area of the individual light sub-regions recovered by connected-component labelling of the light-classified pixels within the object.

